# The bacterial quorum sensing signal DSF hijacks *Arabidopsis thaliana* sterol biosynthesis to suppress plant innate immunity

**DOI:** 10.1101/2020.01.30.927731

**Authors:** Tuan Minh Tran, Zhiming Ma, Alexander Triebl, Sangeeta Nath, Yingying Cheng, Ben-Qiang Gong, Xiao Han, Junqi Wang, Jian-Feng Li, Markus R. Wenk, Federico Torta, Satyajit Mayor, Liang Yang, Yansong Miao

**Affiliations:** School of Biological Sciences, Nanyang Technological University, Singapore 637551, Singapore; Singapore Lipidomics Incubator (SLING), Department of Biochemistry, YLL School of Medicine, National University of Singapore, Singapore, Singapore; Institute for Stem Cell Biology and Regenerative Medicine, Bellary Road, Bangalore, 560065, India; Manipal Institute of Regenerative Medicine, Manipal Academy of Higher Education, Bangalore, 560065, India; Singapore Centre for Environmental Life Sciences Engineering, Nanyang Technological University, Singapore 637551; School of Life Sciences, Sun Yat-sen University, Guangzhou 510275, China; Department of Biology, Southern University of Science and Technology, Shenzhen, China, 518055; National Centre for Biological Sciences, Tata Institute for Fundamental Research, Bellary Road, Bangalore 560065, India; School of Medicine, Southern University of Science and Technology, Nanshan District, Shenzhen, China, 518055; School of Chemical and Biomedical Engineering, Nanyang Technological University, Singapore 637459, Singapore

**Author notes:** These authors contributed equally (Tuan Minh Tran, Zhiming Ma, Alexander Triebl, Sangeeta Nath). Syngenta, Münchwilen AG, Canton of Aargau, Switzerland. For correspondence: Yansong Miao.

**Keywords:** quorum sensing, plasma membrane, endocytosis, plant-pathogen interaction

## Abstract

Quorum sensing (QS) is a recognized phenomenon that is crucial for regulating population-related behaviors in bacteria. However, the direct specific effect of QS molecules on host biology is largely under-studied. In this work, we show that the QS molecule DSF (*cis*-11-methyl-dodecenoic acid) produced by *Xanthomonas campestris* pv. *campestris* can suppress pathogen-associated molecular pattern (PAMP)-triggered immunity (PTI) in *Arabidopsis thaliana*, mediated by flagellin-induced activation of flagellin receptor FLS2. The DSF-mediated attenuation of innate immunity results from the alteration of oligomerization states and endocytic internalization of plasma membrane FLS2. DSF altered the lipid profile of *Arabidopsis*, with a particular increase of the phytosterol species, which impairs the general endocytosis pathway mediated by clathrin and FLS2 nano-clustering on the plasma membrane. The DSF effect on receptor dynamics and host immune responses could be entirely reversed by sterol removal. Together, our results highlighted the importance of sterol homeostasis to plasma membrane organization and demonstrate a novel mechanism by which pathogenic bacteria use their communicating molecule to manipulate PAMP-triggered host immunity.

**SIGNIFICANCE STATEMENT:** Bacteria rely on small signalling molecules called quorum sensing (QS) signals to communicate and coordinate their behaviors. QS is known to regulate gene expression, production of virulence factors, and biofilm formation for pathogenic bacteria to effectively colonize their hosts and cause diseases. In this work, we found a class of QS molecule called diffusible-signal factor (DSF), produced by devastating phytopathogenic bacteria such as *Xanthomonas* spp. and *Xylella fastidiosa*, could communicate directly with plant host and subvert plant innate immunity by inducing plant sterol production and thereby, attenuating receptor signalling through hindering the receptor clustering and plant endocytosis. The results significantly enrich our understanding of the mechanisms in the tug-of-war between bacterial pathogenesis and host immunity.

## INTRODUCTION

Bacteria use quorum sensing (QS) to precisely coordinate population behaviors in response to various environmental cues. QS signals are small molecules that contribute to bacterial virulence in bacterial-host interactions by either regulating bacterial type III secretion or priming host immune systems (1–3). Although the contribution of quorum sensing signals to bacterial pathogenicity has been examined extensively, the direct effects of these small molecules on host biology remain under-explored.

QS signals produced by human pathogens, such as N-3-oxododecanoyl homoserine lactone (3OC12-HSL), modulate pathogen-associated molecular patterns (PAMPs)-mediated NF-kB signaling in macrophages and innate immune responses in recognition of LPS — thereby promoting persistent infection (4). The 3OC12-HSL was also shown to be recognized by the bitter taste receptor, T2R family protein, T2R38 in neutrophils, lymphocytes and monocytes to activate innate immune responses (5). Similarly, plant pathogenic bacteria also secrete small-molecule virulence factors to cross-talk with host and manipulate host immunity during infection. QS molecules produced by several bacterial phytopathogens showed a wide range of interference in host development and defence mechanisms, including the root morphological development in an auxin-dependent manner or callose deposition during immune responses (6–8). However, the detailed mechanisms of how QS molecules directly influence plant host development and pathology, especially the PAMP-mediated host immunity, remains elusive.

Here, we studied the host-pathogen communication between host *A. thaliana* and a specific QS molecule, the diffusible signal factor (DSF). The DSF QS molecule family is produced by diverse Gram-negative bacteria, many of which are devastating phytopathogens such as *Xanthomonas campestris* pv. *campestris (Xcc)*, *Xanthomonas oryzae* pv. *oryzae*, *Xylella fastidiosa*, as well as several human pathogens, including *Pseudomonas aeruginosa, Stenotrophomonas maltophilia*, and *Burkholderia cenocepacia* (9). Notably, *Xanthomonas campestris pv. campestris (Xcc)*, a global-threat phytopathogen that causes black rot on crucifers (10), produces up to four different analogs of DSF (*cis*-2 unsaturated fatty acids), with DSF(*cis*-11-methyl-dodecenoic acid) being the primary compound (>75%) at concentrations of 40-100 μM during infection (11). DSF regulates the expression of *Xcc* Type III secretion system, host cell-wall macerating enzymes and other traits *in Xcc* (9, 12), and elicited host immune response associated with an enhanced *Arabidopsis PR-1* gene expression (11). However, the mechanism by which this QS molecule modulates plant defense responses is mostly unknown.

In this study, we investigated the mechanisms by which DSF alters host physiology and pathology. We demonstrated that DSF induces multiple macro- and microscopic changes in *A. thaliana* cell biology and development, including root morphogenesis and plant lipid profiles. The DSF-induced phytosterols production that impaired both clathrin-mediated-endocytosis (CME) for internalizing FLS2 and FLS2 nano-clustering on the cell surface, which therefore desensitized *Arabidopsis* immune responses to the bacterial flagellum. Removing the DSF-induced sterol accumulation in *Arabidopsis* could reverse the defects in host endocytosis, development, and immunity caused by DSF. However, the FLS2-BAK1 interaction and MAPK activation by flg22 elicitation were unchanged, indicating uncoupled PTI responses underlying DSF immune-signalling. Our studies have significant implications for the understanding of bacterial QS molecule DSF function in host biology and host PAMP-triggered immunity (PTI) pathways.

## RESULTS

### DSF suppresses bacterial flagellin-triggered innate immune responses of *Arabidopsis*

Plants have developed multiple sophisticated defense mechanisms to recognize and respond to a wide range of bacterial virulence factors. Moreover, bacterial pathogens strategically instigate or subvert the host immune response by interfering with PAMP recognition (13, 14). We asked whether DSF, a recently discovered QS signal produced by diverse Gram-negative pathogens (15), could dysregulate pattern-triggered immunity (PTI) responses. We first examined *Arabidopsis* growth in response to DSF molecules within the pathological concentration of DSF during infection (11) and found that DSF inhibited primary root growth of *A. thaliana* in a dose-dependent manner without apparent effects on seed germination (Fig. S1A). To test whether DSF influences flagellin-triggered immunity, we tested several flagellin-triggered *Arabidopsis* responses in plants treated with 25 μM of DSF, the concentration at which root growth was inhibited by 40%. We found that flg22-induced stomatal closure (aperture reduced from 4 μm to 2 μm) was impaired by DSF treatment (Fig. 1A). Callose deposition, a plant physical defense responses to prevent pathogens infections (16), was also significantly impaired in DSF-treated seedlings (Fig. 1B) as was flg22-induced Reactive Oxygen Species (ROS) burst by half (Fig. 1C). We next performed a plant infection assay, in which *Arabidopsis* plants pre-treated with DSF or DMSO (mock) were inoculated with *Pseudomonas syringae pv. tomato* DC3000 (*Pst* DC3000). The use of a non-DSF-producing bacterium here (*Pst* DC3000) was to differentiate the direct effect of DSF on plant immunity vs. the effect of DSF through the activation of the bacterial Type III secretion system and other traits regulated by QS, which would otherwise be difficult to dissect as DSF-deficient or overproducing mutants are less virulent on plants. In this assay which involved using live bacteria with all the potential PAMPs, we found that DSF-treated plants were colonized by a higher number of *Pst* DC3000 cells compared to control plants, indicating a higher susceptibility of plants to *Pst* infection in the presence of DSF (Fig. 1D). We further examine the ability of this QS molecule in compromising the defense mechanism of the plants that were pre-protected by PTI-signaling. In treatments where plants were primed with flg22 peptide before bacterial inoculation, we found that DSF application prior to flg22 priming significantly lowered flg22-induced protection against infection, reflected in a higher bacterial population in DSF+flg22 treatment compared to that of DMSO+flg22 treatment (Fig. 1D-E).

**Fig. 1.**
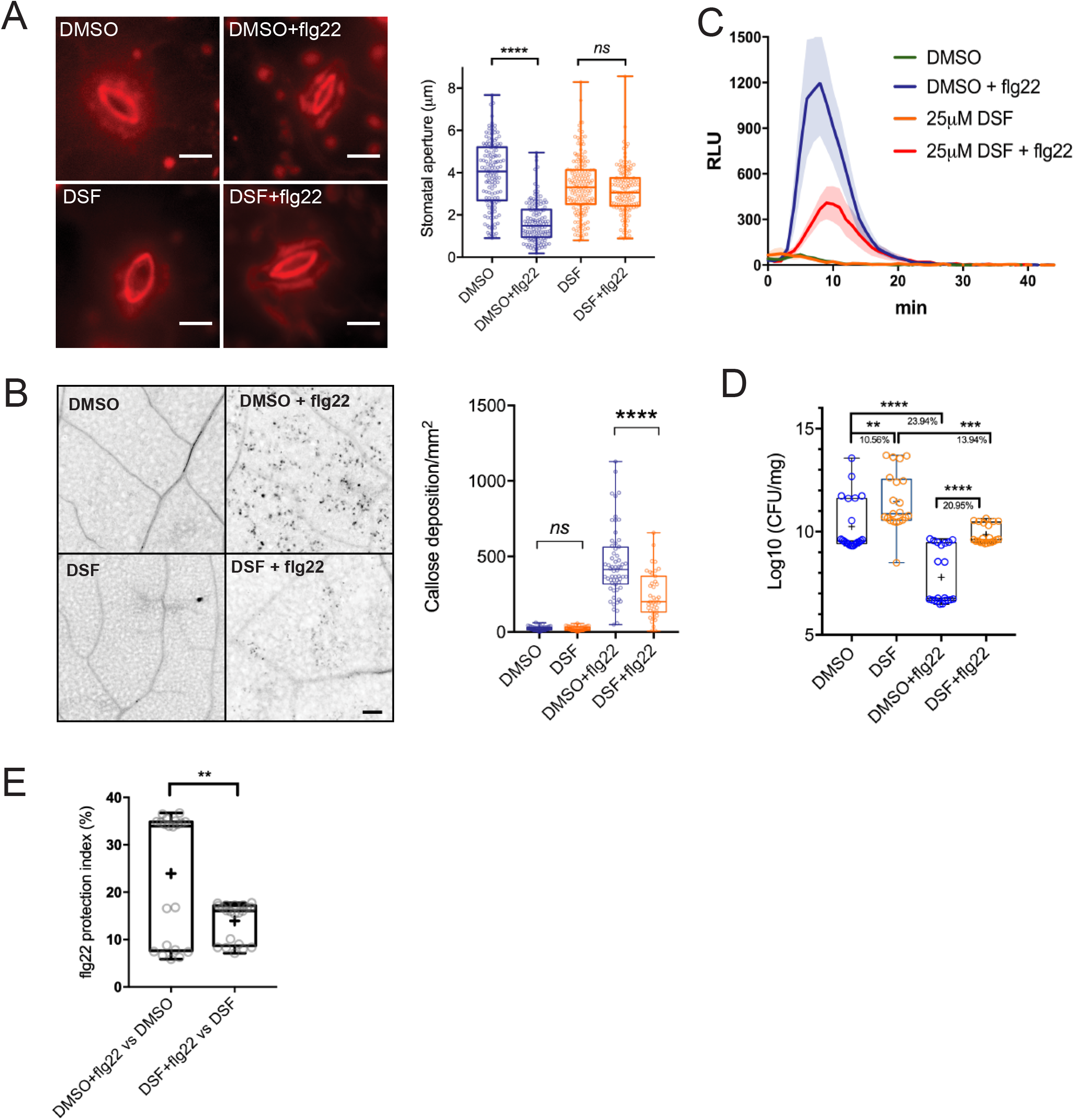
Diffusible-signal factor (DSF) simultaneously dampens several plant defense responses in *Arabidopsis thaliana* against the bacterial PAMP flg22. **(A)** Stomatal apertures of 5-week old Col-0 leaves after flg22 elicitation. Intact leaves were infiltrated with DMSO or 25 μM DSF diluted in 10 mM MgCl_2_ for 24 h before the epidermal layers of detached leaves were peeled off and treated with 1 μM flg22, stained with propidium iodine and imaged. Stomatal apertures were measure by Fiji (n≥130 stomata, bars, 10 μm). **(B)** Callose deposition in 2-week old Col-0 plants pre-treated with DMSO or 25 μM DSF in ½ liquid MS for 24 h before being elicited by flg22 in liquid ½ MS. Leaves were stained with aniline blue and imaged using a confocal microscope with UV excitation to visualize flg22-triggered callose deposition. Plants treated with only DMSO or DSF were used as negative control (n≥40 images; bar, 100 μm) **(C)** Reactive oxygen burst in Col-0 leave strips pre-treated with DMSO or 25 μM DSF for 24 h prior to flg22 elicitation (n≥6 leaf discs/condition). The experiment were repeated three times. **(D)** Bacterial population at 4 days post-inoculation (DPI) in *Arabidopsis* 2-week old seedlings pre-treated with DMSO (solvent control) or DSF, followed by priming with flg22 peptide and flooding assay with *Pst* DC3000. Boxplot represent pooled data from two independent experiments (n=21). **(E)** Protection index (%) was derived from log CFU/mg tissue by dividing the differences between flg22-treated group and the control group to the mean bacterial population of the control group. P-values were determined by one-way ANOVA (^∗^ p < 0.05; ^∗∗^ p < 0.01; ^∗∗∗^ p < 0.001; ^∗∗∗∗^ p < 0.0001; ns, not significant).

### DSF interferes with the host endocytic internalization on the plasma membrane without compromising MAPK signaling

We next sought to investigate if DSF could interfere with the dynamics and functions of the *Arabidopsis* flagellin receptor FLS2 (16). Flagellin binding leads to conformational changes in the FLS2 receptor and triggers its oligomerization (17) and endocytic internalization to activate the innate immunity cascade (18, 19).

We first tested whether DSF affects FLS2 endocytosis upon elicitation with its corresponding ligand flg22. Upon treatment of flagellin peptide flg22 but not flgII-28, FLS2 endocytic internalisation was evidenced by the appearance of punctate endosomes after around 60-75 min and replenishment of FLS2-GFP back to the plasma membrane (PM) at 120 min post elicitation (Fig. 2A and Fig. S1B) (18, 20, 21). In contrast, when FLS2-GFP seedlings were treated with DSF for 24 h prior to flg22 exposure, we observed an apparent delay in FLS2 internalization upon PAMP elicitation. After 60 mins of flg22 elicitation, DSF-treated plants showed fewer FLS2-positive endosomes compared to DMSO-treated plants, indicating the attenuated endocytosis of FLS2. We then asked whether the attenuated FLS2 endocytosis could also lead to a delay in FLS2 degradation (22, 23). Consistent with a previous observation (24), we found that FLS2 degradation occurred approximately 30 min after peptide elicitation and this response was similarly delayed in DSF pre-treated seedlings (Fig. S1C).

**Fig 2.**
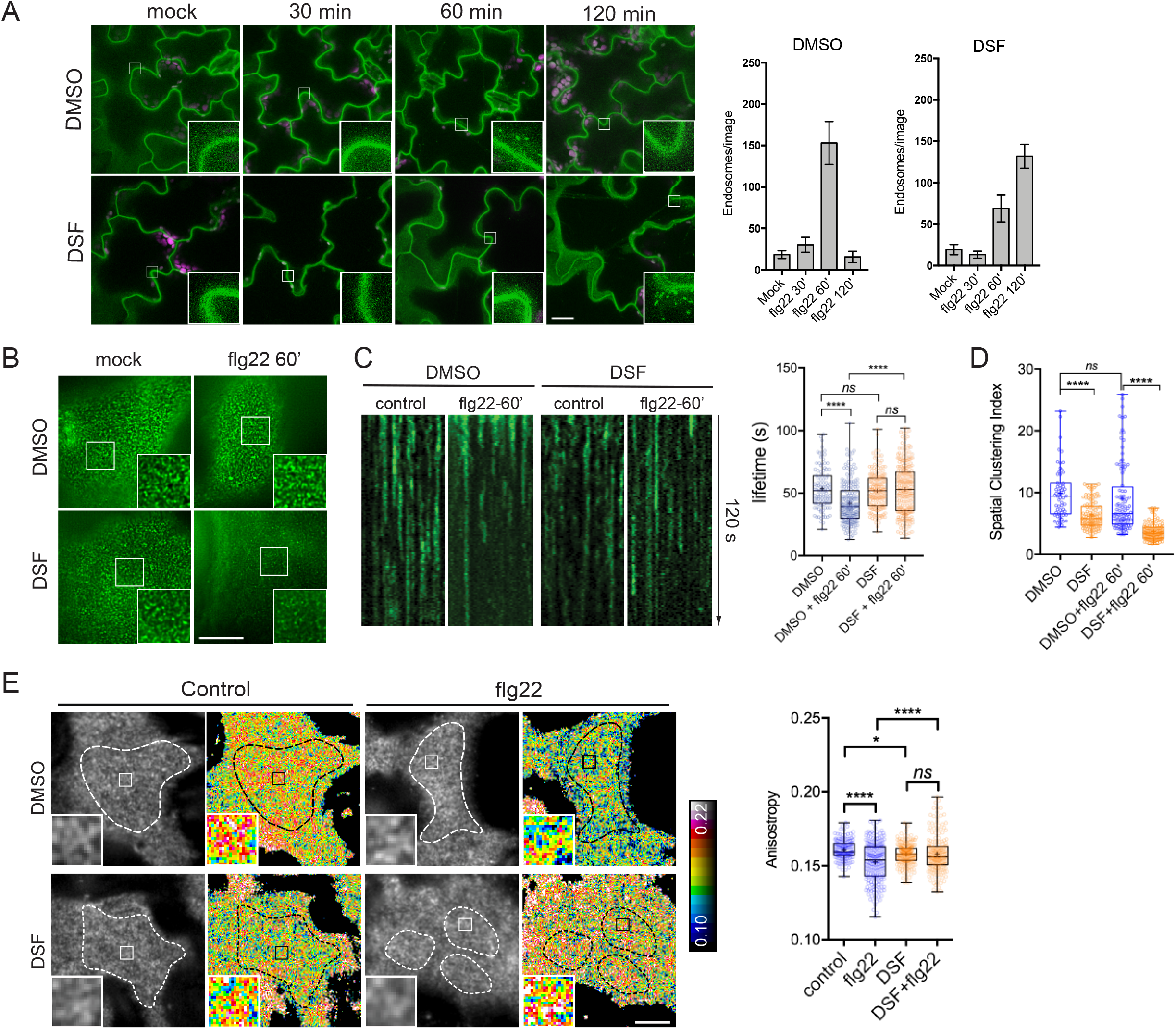
DSF delays ligand-induced endocytosis of the FLS2 receptor in response to flg22 peptide. **(A)** Five-day old FLS2-GFP seedlings were treated for 24 h in liquid 1/2 MS medium supplemented with 25 μM DSF or DMSO (solvent control) before being elicited with 10 μM flg22 and imaged at indicated timepoints. Micrographs represent maximum projections of 12 slices every 1 μm z-distance (bars, 20 μm). Bar graphs represent the numbers of endosomes per image. Error bars, SEM (n≥12) **(B)** FLS2-GFP receptors on the PM of *Arabidopsis* cotyledons. Five-day old FLS2-GFP plants were treated with DMSO or DSF for 24 h prior to elicitation by 10 μM flg22 and imaged using VA-TIRFM (bar, 1μm). **(C)** Kymographs of FLS2-GFP clusters on *Arabidopsis* plasma membrane before and after 60 min of flg22 elicitation, following DMSO or 25 μM DSF 24-h pre-treatment. The lifetime of FLS2-GFP foci was measured from kymographs using Fiji (n≥ 100 endocytic events). **(D)** Boxplot represents spatial clustering of FLS2 receptor foci, indicated by Spatial Clustering Index (SCI). SCI was calculated from TIRF timelapse movies based on the ratio of top 5% of pixels with the highest intensity and 5% of pixels with the lowest intensity (n≥60 ROIs). **(E)** Homo-FRET analysis of receptor oligomerization on PM of DMSO- and DSF-treated FLS2-GFP seedlings upon flg22 elicitation for 5 min. As the plasma membrane is not definite flat, anisotropy quantification was performed only in those well-focused regions of the plasma membrane (white dashed lines) (bar, 5 μm). Boxplot represents anisotropy values calculated from homo-FRET imaging of FLS2-GFP. Representative intensity and anisotropy images were shown. Insets are representative 20×20 pixel ROIs used for data analysis (n≥150 ROIs). P-values were determined by one-way ANOVA (^∗^ p < 0.05; ^∗∗^ p < 0.01; ^∗∗∗^ p < 0.001; ^∗∗∗∗^ p < 0.0001; ns, not significant).

To quantitatively determine the DSF-caused defects in receptor endocytosis, we examined the FLS2 endocytic internalization at the plasma membrane (PM) using variable-angle epifluorescence microscopy (VAEM) (Fig. 2B). From VAEM micrographs, we quantitatively determined receptor internalisation by kymograph analysis of FLS2-GFP puncta. We observed that flg22 enhanced FLS2 endocytosis as shown by the shortened lifetime of FLS2 on the plasma membrane, and this enhancement was blocked by DSF pre-treatment prior to peptide elicitation (Fig. 2C). As FLS2 is endocytosed via clathrin-mediated endocytosis (CME) (20), we then tested whether the delay in FLS2 endocytosis was due to a general inhibition of CME. Indeed, we observed a significant increase in the lifetime of clathrin-light-chain (Fig. S2A), as well as the lifetime of several typical CME-dependent endocytic receptor: brassinosteroid receptor BRI1 and Boron receptor BOR1 under DSF treatment (Fig. S2B-C), indicating the impaired endocytic internalization of these PM receptors.

Unexpectedly, although DSF delayed FLS2 degradation and recycling due to the impaired CME (Fig. S1C), we did not observe a noticeable change in the acute response of MAPK phosphorylation, which occurs approximately 15 min post-elicitation and returns to basal level within 60 min (Fig. S1D). This result suggests that flg22-induced MAPK activation is independent of FLS2 endocytosis under DSF-signalling, which is consistent with the unchanged *FRK1* up-regulation in the presence of flg22 (Fig. S1E) that is through the activation of a MAPK signalling cascade (25). Exposure to flg22 could still induce the expression of *FRK1* in DSF-treated seedlings, suggesting that DSF did not have a significant effect on *FRK1*-related pathway and might not influence FLS2-BAK1 heterodimerization that was critical for *FRK1* induction (26). In agreement with this observation, we also found that DSF did not affect the flg22-induced FLS2-BAK1 heterodimerization (Fig. S1F), suggesting that the inhibition of flg22-induced ROS burst by DSF is unlikely a direct consequence of disturbed FLS2-BAK1 complex formation, and uncoupled from MAPK activation.

### DSF impairs lateral nano-clustering of FLS2 on the plasma membrane

In addition to the endocytic internalization that removes material from the plasma membrane, the lateral nano-clustering of plasma membrane-bound receptors through multivalent interactions is well-known to mediate diverse signal transduction mechanisms during host-pathogen interactions (27; 28). The self- or hetero-oligomerization of surface receptors are critical for receptor activation, signal amplification, and signal transduction in both plants and human, such as the brassinosteroid receptor BRI1 (29), chitin receptor *At*CERK1 (30), EGF receptor (31, 32) or insulin receptor IR/IGFR (33). To further investigate DSF-induced effects on receptor lateral dynamics and their activity, we performed living cell imaging of FLS2-GFP, chosen from several DSF hindered endocytic PM receptors (Fig. 2C, Fig. S2B-C). FLS2 formed heterogeneous PM clusters with or without ligand activation (Fig. 2B), representing the resting- or activated-states, respectively (34). The formation of FLS2-BAK1 heterodimer upon flg22 elicitation is a key step in activating downstream defense signaling (35, 36). Besides the FLS2-BAK1 interaction, an increase in FLS2-FLS2 self-oligomerization is also important for FLS2 phosphorylation and signal transduction (37). Though DSF did not influence flg22-triggered FLS2-BAK1 interaction and MAPK activation (Fig. S1D,F), the fact that DSF impaired the ROS production suggested uncoupled mechanisms in the activation of MAPK and the ROS production by potential using differential inter- or intra-protein interactions of FLS2 for these processes. Consistently, with a DSF pre-treatment, FLS2 clusters became more diffused with a lower spatial-clustering index (SCI) after 60 min of flg22 elicitation (Fig. 2D).

To study the detailed physical interactions of FLS2 receptors in the multivalent nanoclusters, we took advantage of a well-established living cell imaging approaches in studying mammalian receptor clustering and activation (38, 39). We quantitatively measured the lateral self-interaction of FLS2 on the plasma membrane directly under ligand activation by performing FRET between identical fluorophores (homo-FRET) coupled with VAEM, which allows sensitive measurement of fast cargo-induced receptor activation (40). In this method, homo-oligomerization of FLS2-GFP will be accompanied by a reduction of fluorescence emission anisotropy of FSL2-GFP when it is excited by polarized light. We determined the steady-state fluorescence emission fluorescence anisotropy of FLS2-GFP with or without 5 minutes of peptide elicitation (Fig. 2E, Fig. S2D,E). Flg22 stimulated a significant decrease in anisotropy of FLS2-GFP, reflecting the flg22-induced oligomerization of FLS2. However, DSF-treated plants showed a reduction of flg22-triggered anisotropy change, indicating a suppressed formation of homo-oligomer (active) from homo-dimer states (unstimulated) (41). Seedlings exposed to DSF did not exhibit any significant change in emission anisotropy of FLS2 after flg22 elicitation (compared to DMSO+flg22 control, p>0.05) (Fig. 2E), consistent with SCI analysis (Fig. 2D).

We next investigated the effects of DSF on bulk endocytosis using the lipophilic dye FM4-64 and CME using transgenic plant expressing PIN2-GFP (42). Briefly, we monitored the uptake of FM4-64 in *Arabidopsis* seedlings expressing PIN2-GFP, an auxin efflux carrier protein and a CME-dependent cargo (43) in the presence of the novel small-molecule inhibitor ES9-17, which specifically binds to clathrin heavy-chain without protonophoric effect (44, 45). To test whether DSF would influence CME further on top of a partially-compromised CME by small molecular inhibitors (46), we pre-treated PIN2-GFP seedlings with DMSO/DSF and then, performed pharmacological studies with different combinations of ES9-17/BFA on treated plants, using FM4-64 dye as an indicator of bulk endocytosis. Our result indicated that while neither DSF nor 50 µM ES9-17 completely blocked the internalization of FM4-64 dye, a combined treatment of DSF and ES9-17 totally inhibited FM4-64 uptake, indicated by the absence of intracellular FM4-64 signal and FM4-64-containing BFA bodies in BFA treatment (Fig. 3A-D). Our results demonstrated the additional attenuation of bulk-phase endocytosis by DSF on top of the CME inhibitor ES9-17.

**Fig. 3.**
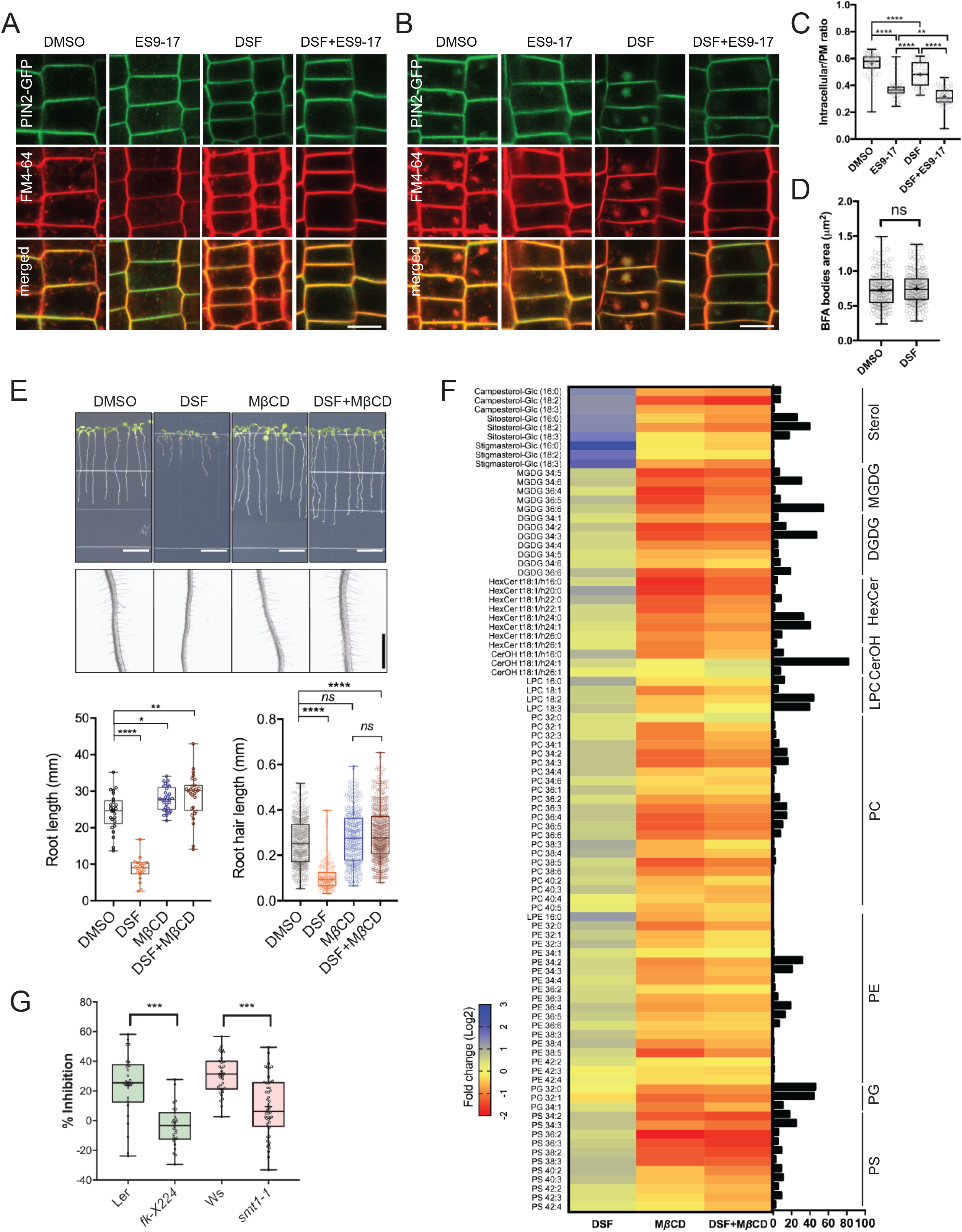
DSF inhibited endocytosis in *Arabidopsis* roots by altering plant lipid profile. **(A)** Internalization of the endocytic tracer FM4-64 in PIN2-GFP seedling roots pre-treated with DMSO or DSF for 24 h. Plants were treated in liquid ½ MS with DMSO (control) or 50 μM ES9-17 for 1 h before being stained with FM4-64 and visualised using confocal microscopy (n≥5 plants; bars, 10 μm). **(B)** Accumulation of intracellular FM4-64 signals into BFA compartments after BFA treatment. PIN2-GFP seedlings pre-treated for 24 h on ½ MS agar supplemented with DMSO or DSF were subjected to BFA or ES9-17+BFA treatments before being stained with FM4-64 and imaged (n≥5 plants; bars, 10 μm). (**C**) Boxplot represents the ratio of intracellular/plasma membrane signal intensity of FM4-64 dye in root cells presented in Fig. 3A as an indicator of bulk endocytosis (n≥37 cells). (**D**) Boxplot represents the area of BFA compartments in seedlings grown on DMSO- or DSF-supplemented plates as presented in Fig. **3B**, n≥161 BFA compartments. **(E)** Growth of Col-0 seedlings on 1/2 MS agar supplemented with DMSO, 25 μM DSF, 2mM M*β*CD and 25 μM DSF+2mM M*β*CD. Images of whole seedlings and root hairs were taken after seven days (top panel bars, 10 mm; bottom panel bar, 1mm). The length of primary roots (n≥20) and root hairs (n≥140) of 7-day old seedlings on different medium measured by Fiji. **(F)** Lipidomic profiles of *Arabidopsis* Col-0 seedlings after seven days growing on 1/2 MS agar supplemented with either 25 μM DSF, 2 mM M*β*CD or 25 μM DSF + 2 mM M*β*CD. Heatmap representing the log2 value of fold change compared to DMSO treatment (n=6 replicates). The bar graph parallel to the heatmap indicates the relative abundance (in percentages) of the corresponding species within each lipid class. **(G)** DSF-induced Inhibition rate of sterol mutants *fk-X224* and *smt1-1* and their corresponding wildtype ecotypes Ler and Ws. Seedlings were monitored for seven days on DMSO- and 25 μM DSF-supplemented ½ MS plates (n≥28 plants). Inhibition rate (%) was represented by the ratio between the difference in primary root length of seedlings on control (DMSO) vs. DSF-supplemented medium and the mean primary root length of control seedlings. P-values were determined by one-way ANOVA (^∗^ p < 0.05; ^∗∗^ p < 0.01; ^∗∗∗^ p < 0.001; ^∗∗∗∗^ p < 0.0001; ns, not significant).

We then asked whether such impairment of plant CME could also disrupt endosomal transport after endocytic invagination from the plasma membrane. We examined the intracellular membrane compartments of the protein sorting and endosomal transport pathways upon DSF treatment. No noticeable ultrastructural changes were observed in early endosome/trans-Golgi network (VHA-a1), late endosome/multivesicular body (MVB) (RabF2b), and Golgi (Got1p homolog), nor in their response to Brefeldin A (BFA) or Wortmannin that induce Golgi aggregation or MVB dilation (47), respectively (Fig. S3A-D). The above results suggest that DSF likely impairs plant endocytosis mainly at the step of endocytic internalization on the plasma membrane and not at the endomembrane trafficking steps.

### DSF increased host phytosterol contents and reprogramed *Arabidopsis* lipid metabolism

Dynamic behaviors of PM proteins, including the endocytic invagination, lateral diffusion, and inter-molecular interactions, rely on the lipid-compartmentalization and a balanced sterol composition of the plasma membrane (39, 48). We next sought to determine if the lipid profile of *Arabidopsis* was also altered by DSF, given that this QS signal is a short-chain fatty acid-like molecule that could serve as a building block to create diverse biologically active molecules (2). Our targeted lipidomic profiling of *Arabidopsis* seedlings revealed that *Arabidopsis* seedlings grown on DSF-supplemented medium had a comprehensive shift in a wide range of lipid compounds compared to those grown on control medium (Fig. 3E-F; Dataset S1-2). Interestingly, the most notable changes were observed in several acyl steryl glycosides (ASGs), of which the most abundant phytosterols: sitosterol, campesterol, and stigmasterol showed an increase by 2-4 folds. Sterols regulate a wide range of cellular processes, such as lipid metabolism and cell membrane dynamics, such as the formation of membrane microdomains (49, 50). We, therefore, tested whether the increased sterol content could explain the DSF-induced phenotypes by treating *Arabidopsis* seedlings with Methyl-*β-*cyclodextrin (M*β*CD) in combination with DSF. M*β*CD is a widely-used sterol-depleting compound that acts strictly at the membrane surface by binding to sterol with high-affinity and remove sterol acutely within minutes, with demonstrated success in acute depletion of sterols in both plant and animal cells (51–55). Here, we found that M*β*CD fully blocked the DSF-induced inhibition of primary roots and root hairs, suggesting DSF’s opposite effects compared to M*β*CD on sterol perturbation in *Arabidopsis*. Consistent with this result, culturing seedlings in the presence of both M*β*CD and DSF completely abolished the DSF-triggered increase of lipid components, including phytosterols. The total quantity of measured sterols content in M*β*CD and M*β*CD+DSF treatment were reduced to 92.4 ± 10.3 nmol.g^−1^ and 76.0 ± 10.4 nmol.g^−1^ respectively, compared to 138.2 ± 24.6 nmol.g^−1^ in DMSO treatment and 366.3 ± 121.1 nmol.g^−1^ in DSF treatment, showing a dominant effect of M*β*CD over DSF (Fig. 3F).

To further examined the role of phytosterols in DSF-induced inhibition of primary root growth, we performed growth assay on several *Arabidopsis* mutants of the sterol biosynthesis pathway (Fig. 3G) in the presence of DSF. The *smt1-1* and *fk-X224* mutants were chosen for this experiment as their growth defects are less severe compared to other sterol mutants. Primary root growth of *smt1-1* mutant (defective in C-24 methyltransferase) and the *fackel (fk)* mutant *fk-X224* (defective in sterol C-14 reductase), which are two key early steps of sterol biosynthetic pathway upstream of the branch point (56) (Fig. S3E), showed significantly reduced sensitivity to DSF, as represented by a lower inhibition rate compared to their corresponding wild-type ecotypes (Fig. 3G, Fig. S3F). Together, these findings support our hypothesis that DSF regulates host development through altering host phytosterols content.

### Sterols regulation of lipid microdomain assembly and endocytosis phenocopies DSF function in *Arabidopsis*

As we observed that M*β*CD and DSF have opposite effects on *Arabidopsis* lipidomic profiles in the long-term plant growth assay, we next examined whether an acute sterol removal by M*β*CD would restore other DSF-induced phenotypes on plant endocytosis and innate immune responses of *Arabidopsis*. In the experiments testing DSF perturbation of PTI, following M*β*CD pre-treatment, the number of FLS2-positive endosomes in FLS2-GFP plants pre-treated with DSF increased significantly at 60 min after elicitation with flg22, even though M*β*CD itself strongly inhibited FLS2 trafficking into endosomes (Fig. 4A). In line with this finding, M*β*CD also restored the bulk endocytosis of FM4-64 dye that was blocked by the combined treatment of DSF and ES9-17, represented by the similar ratio of intracellular/PM signal intensity and BFA bodies area between DMSO and ES9-17 treatment in the DSF+M*β*CD pre-treated seedlings (Fig. 4B and Fig. S4A), the delay of CLC lifetime (Fig. S4B), and the DSF-induced inhibition of ROS burst (Fig. 4C).

**Fig. 4.**
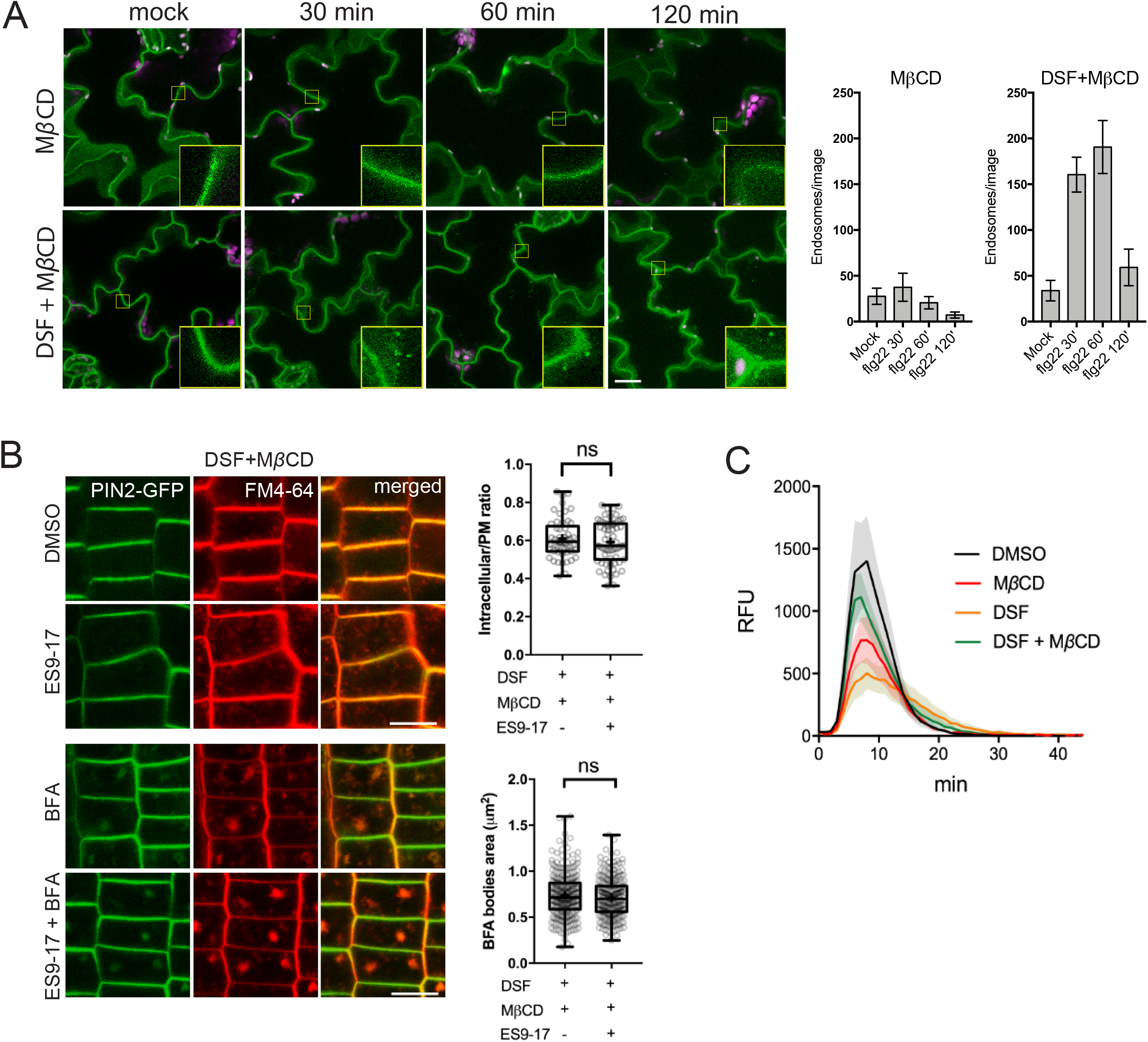
Sterol depletion by M*β*CD recovered DSF/sterol-induced phenotypes on *Arabidopsis* seedlings. **(A)** FLS2 receptor internalization into endosomes monitored by confocal microscopy following 10 µM flg22 treatment. FLS2-GFP seedlings were pre-treated for 24 h with DMSO or 25 μM DSF in ½ MS, followed by a brief 2 mM M*β*CD treatment for 30 min prior to peptide elicitation. Micrographs represent maximum projections from 12 slices taken every 1 μm z-distance. Bar graphs showed the number of endosomes per image (n≥10 images; bar, 20 μm). (**B**) Internalization of FM4-64 dye of PIN2-GFP seedling roots after plants were treated for 24 h on ½ MS plates with 25 μM DSF followed by 30 min treatment with 2 mM M*β*CD in liquid ½ MS. Plants were treated with 50 μM ES9-17 for 1 h and/or 50 μM BFA-1h before staining with FM4-64, and DMSO was used as solvent control (n≥5 plants; bar, 10 μm). Quantification of intracellular/PM ratio were performed on n≥50 cells and BFA bodies area measurement were performed on n≥250 BFA bodies. **(C)** ROS production by Col-0 leaf strips treated for 24 h with DMSO or 25 μM DSF, followed by a 30-min treatment with water (control) or 2 mM M*β*CD prior to elicitation by 1 μM flg22 peptide. The experiment were repeated twice with similar result. P-values were determined by 2-tailed Student’s *t*-test (^∗^ p < 0.05; ^∗∗^ p < 0.01; ^∗∗∗^ p < 0.001; ^∗∗∗∗^ p < 0.0001; ns, not significant).

We further asked whether the supplementation of phytosterols could phenocopy the DSF-induced defects in CME and flg22-activated FLS2 internalization. We found that exogenous phytosterol mix (Sitosterol: Stigmasterol: Campesterol=8:1:1) treatment phenocopied DSF-induced inhibition of flg22-stimulated ROS production and subsequent M*β*CD supplement reversed the sterol-inhibited endocytosis of FM4-64 dye in the presence of ES9-17 (Fig. S4C-D), FLS2 internalization into endosomes (Fig. S5A), as well as flg22-triggered ROS burst (Fig. S5B). Similar to the sterol depleting compound M*β*CD, we observed that Lovastatin— a sterol biosynthesis inhibitor— also rescued ROS burst in DSF-treated leaves (Fig. S5C), suggesting that the DSF-induced inhibition of ROS and innate immunity was likely due to the incorporation of DSF into phytosterol metabolic pathways or induction of sterol production via an unknown mechanism. Similar inhibitions of plant root growth were also observed with the predominant sterol *β*-sitosterol: *β*-Sitosterol could also inhibit root growth in a dose-dependent manner similar to DSF, and this inhibition could also be reverted by M*β*CD treatment (Fig. S5D-E). Together, these data suggest that a balance in sterol composition of the plant PM is crucial for plant CME as well as the pattern-recognition receptors (PRRs)-mediated defense responses.

### DSF and Sterol interfere with lateral membrane compartmentalization and FLS2-nanoclustering

Lipids composition, compartmentalization, and surface tension of plasma membrane are critical physicochemical properties in regulating lateral motility, protein-protein interaction, and distribution of immune receptors, including FLS2 (28, 57). Sterols are essential constituents of the liquid-ordered (L_o_) lipid nanodomains and influence membrane flexibility (58). We, therefore, asked whether the DSF-induced phytosterol increase influences the compartmentalization of PM by imaging nanodomain-marker YFP:REM1.2 *Arabidopsis* expressing YFP:REM1.2 treated with DSF, a phytosterol mix (sitosterol: stigmasterol: campesterol=8:1:1) or M*β*CD were examined via super-resolution 2D-Structured Illumination Microscopy (2D-SIM). M*β*CD resulted in a significant reduction of the mean intensity of YFP:REM1.2-marked microdomains, suggesting a reduction in the ordered-phase. The presence of either DSF or exogenous sterol could similarly reverse the M*β*CD effect in lipid compartmentalization (Fig. 5A-B). Furthermore, with M*β*CD post-treatment, DSF no longer attenuated the homo-oligomerisation of FLS2 upon flg22-elicitation, indicated by an anisotropy reduction in living cell homo-FRET imaging (Fig. 5C-D).

**Fig. 5.**
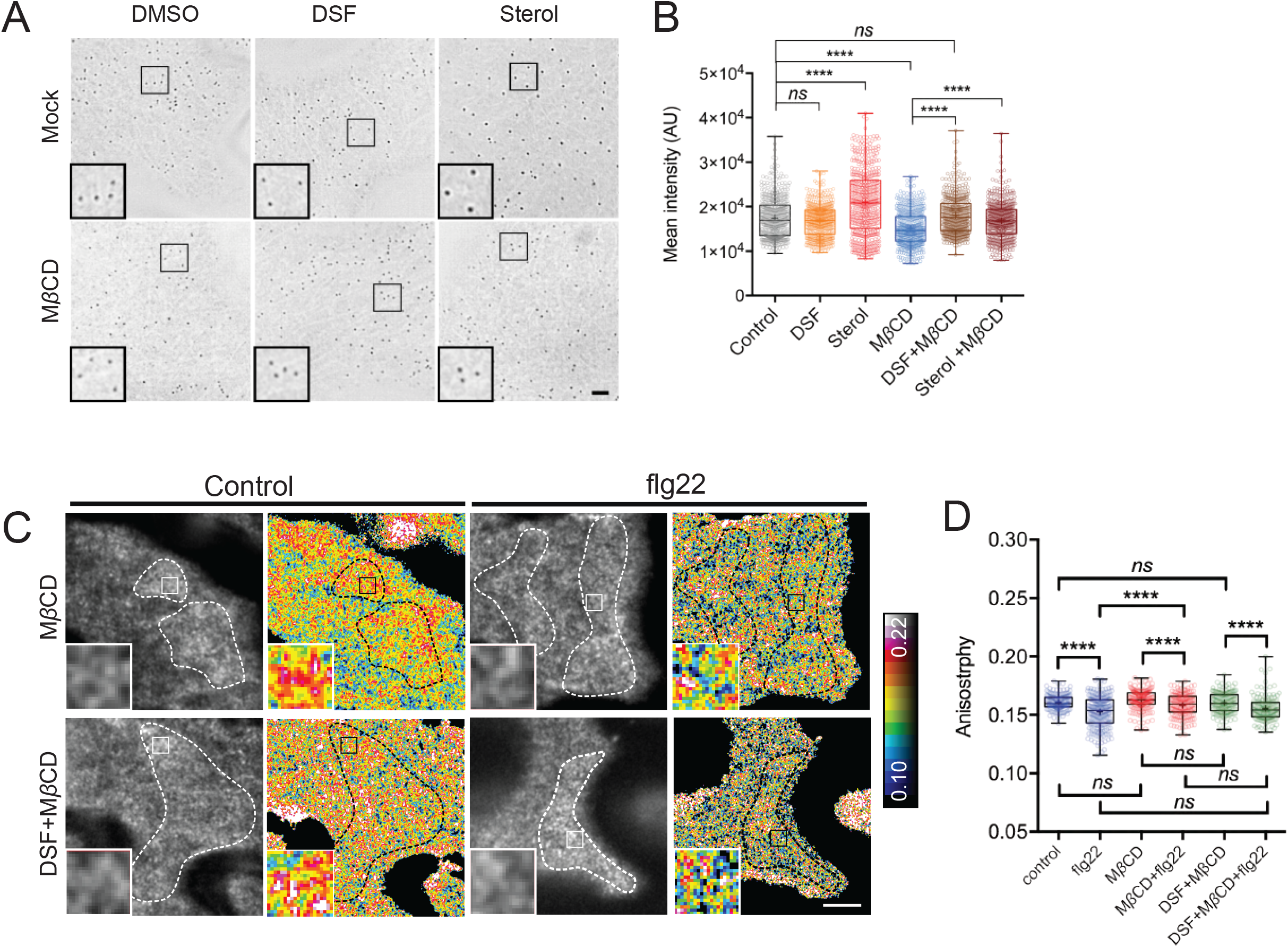
Lipid microdomain integrity is dependent on proper sterol composition of the PM. **(A)** 2D-SIM images of the microdomain marker REM1.2-GFP taken after 24 h treatment with DMSO, DSF, or sterol mix followed by a 30 min treatment with 2 mM M*β*CD or water control (bar, 2 μm). **(B)** Quantification of Remorin foci intensity on PM (n=500 foci). **(C)** Oligomerisation state of FLS2-GFP receptors before and after 5 min of elicitation with 10 μM flg22 following treatment of the seedlings M*β*CD or DSF+ M*β*CD. Representative intensity and anisotropy images were shown. Insets are representative 20×20-pixel ROIs used for data analysis (bar, 5 μm). **(D)** Boxplot represents anisotropy values measured from Homo-FRET experiment (n≥150 ROIs). P-values were determined by one-way ANOVA (^∗^ p < 0.05; ^∗∗^ p < 0.01; ^∗∗∗^ p < 0.001; ^∗∗∗∗^ p < 0.0001; ns, not significant).

### Osmotic stress rescued DSF-induced inhibition of *Arabidopsis* endocytosis

Lipid order increase and microdomain assembly deform the organization of local plasma membrane and thus, alter the line tension of phase-separated PM, which are physical cues in regulating endocytosis and lateral molecular dynamics (48, 59, 60). We were therefore motivated to ask whether osmotic stress, which induces plant PM remodelling (48), could reverse the inhibition of endocytosis caused by DSF. To test if osmotic stress could restore the inhibition of endocytosis and plant growth caused by DSF, we measured primary root length from seedlings grown in combination with different concentrations of osmolytes (NaCl or Mannitol) and DMSO/DSF. We observed a partial recovery of DSF-induced inhibition on primary root growth at 10 and 50 mM NaCl, but not with Mannitol (Fig. S6A). Consistent with this result, flg22-induced internalization of the FLS2 receptor in seedlings, which were incubated with 100 mM NaCl following DSF treatment, could also be observed as early as 30 min after peptide elicitation (Fig. S6B). The differential recovery in root growth by Mannitol and NaCl treatments suggested that while their alteration of PM tension reverts the DSF-attenuated endocytosis similarly, ionic and non-ionic osmolytes may have different mechanisms to achieve a full reversal of root development under DSF-signaling.

## DISCUSSION

Hosts and symbionts constantly exchange small metabolites during interkingdom interactions. For example, the plant pathogenic bacterium *Pseudomonas syringae* produces the phytotoxin coronatine to target multiple plant immunity pathways, including stomatal defence, thus facilitate bacterial entry as well as disease development (61–64). Conversely, the transfer of small molecules from plants to microbes is also an integral part of microbial symbiotic relationships (65, 66).

During host-pathogen interaction, bacteria mainly rely on quorum sensing to regulate the expression of virulence genes necessary for colonization and pathogenesis or to switch between different lifestyles (67–69). Numerous reports highlighted the effects of QS molecules on plant development and immunity, but mechanistic insights on how bacterial QS molecules specifically interfere with host biology remain sparse (70). Distinctive acylation and hydroxylation patterns in bacterial metabolites have been suggested as a molecular signature of AHL QS family to regulate host immune responses (71, 72). For example, medium-chained 3-hydroxy fatty acids directly bind and trigger the phosphorylation of LIPOOLIGOSACCHARIDE-SPECIFIC REDUCED ELICITATION (LORE) receptor kinase, leading to activation of cytoplasmic receptor-like kinases and stimulate *FRK1* expression (71, 73). Here, we unravelled a different mechanism by which a unique QS molecule called DSF regulates host biology by perturbing host phytosterol homeostasis. Unglycosylated sterols and glycosylated sterols (including steryl glycosides and acyl sterol glycosides) are integral components of the plant plasma membranes, especially in membrane microdomains (50, 74). The increase in host acyl-glycosylated phytosterols by DSF treatment shown in our targeted lipidomic analysis might be a consequence of direct sterol modifications – such as DSF being metabolized to a precursor of acyl source for steryl glucoside acyltransferase (75), or from a potential sterol exchange between PM and ER through PM-ER contact sites (76, 77) — both of which require further investigation. Through the modulation of sterols, DSF interferes with plant PTI through the alteration of both the clustering and endocytic internalization of the surface receptors. Compared to untreated plants, DSF-treated plants display an apparent increase in total phytosterol by >2.5 fold, as well as slight enhancement in several other lipid species that are PM structural constituents, such as glycerolipids (PC, PE, PG, and PS) and sphingolipids (ceramides) (78–80). Phosphatidylserine (PS) is known to play an essential function in surface membrane compartmentalization and protein clustering (81, 82). Thus, how the changes of phytosterols and PS by DSF tune the PM mechanical properties is worth of future studies via artificial lipid bilayer-based assays and membrane modelling.

The diverse roles of QS molecules from pathogenic bacteria in both regulating bacterial responses and modulating host biology reflect a direct co-evolution between hosts and pathogens according to the Zig-zag model (83). *Xcc* DSF gradually accumulates during infection and can reach substantial concentrations (hundred micromolar amount) at later stages of disease (11), similar to QS molecules from other bacteria (84). A previous report demonstrated that at high concentrations (e.g. 100 μM), DSF acts as a hypersensitive reaction (HR) elicitor evident by cell-death response and excessive callose deposition (11). Here, we found that at low concentrations (e.g. 25 μM), which is more relevant to the early stages of infection, this QS signal seems to regulate a specific branch of plant PTI responses through the modulation of molecular dynamics of PRRs on the PM. These findings of plant responses to low dose and the high dose of DSF treatments are not necessarily conflicting to each other as they represent different stages of plant immune responses over the accumulation of bacterial DSF during disease development. In addition, the high-dose DSF-induced plant HR could also be masked by bacterial extracellular polysaccharide during plant colonization, demonstrating the constant bidirectional interaction between plant and bacteria (11). Taken together, these above results reflect the dynamic change in balancing the tug-of-war over the accumulation of DSF molecules during bacterial infection. Although we found that DSF did not directly alter PAMP-activated MAPK signaling, heterodimerization of FLS2-BAK1, or *FRK1* expression, this QS molecule did compromise the PAMP-triggered ROS production. Such different DSF-mediated host signalling demonstrates the uncoupled mechanisms of MAPK activation and ROS production upon flg22-elicitation, in which ROS production was attenuated by DSF/sterol while activation of MAPK cascade remained. As a result of a functional MAPK cascade, DSF could only reduce but not entirely abolish the flg22-induced defense responses (Fig. 1D-E). FLS2-BAK1 may not be the only rate-limiting components of the defense signaling cascade as *bak1* null mutants still retain a certain level of MAPK and ROS responses (85), and FLS2-BAK1 association could be uncoupled from MAPK activation as previously shown with the flg22 G/A mutant peptide (26). Similarly, in chitin-mediated PTI, AtCERK1-mediated MAPK-activation and ROS production were also found to be distinctly regulated by receptor-like cytoplasmic kinases (RLCK) PBL27 and BIK1, respectively (86; 87). These observations suggest potential heterogeneity in receptor coupling and distinct constituents in receptor clusters that may convey differential immune-response by recruiting different PTI signaling molecules. Similar scenarios have also been observed in the mammalian innate immune system (88). The FLS2 binding-partners for MAPK and ROS signaling have differential sensitivity to DSF/sterol-induced perturbation of PM, indicating their possibly different affinity or participation in nanodomains and FLS2-nanoclusters during PRRs signaling. Consistently, due to the active involvement in the high-ordered membrane domain of NADPH oxidase proteins, the NADPH oxidase activity has also been reported to be highly sensitive to plant sterol changes (89). The decline of ROS burst under DSF-signaling might also be through the change in signaling competency of other components downstream of FLS2 and BAK1, such as BIK1, BSK1 or heterotrimeric G protein subunits; or the function of the membrane-bound RBOHD itself (90; 91; 92; 93; 94). Of note, the loss-of-function alleles of some of these components have also been documented to affect the flg22-induced ROS burst but not MAPK activation (90; 91; 92; 95).

Increasing evidence supports the roles of membrane microdomains in host innate immunity by regulating surface receptor dynamics in and out of microdomains on plasma membrane upon ligand-recognition, subsequent receptor interactions, and innate immunity activation (27; 96; 97). For instance, disruption of PM continuity is competent enough to abolish the signaling carried out by multiple PM receptors (BRI1, FLS2, EGFR) (28, 98). Similar to many other immune-receptors (99), unstimulated *Arabidopsis* FLS2 could be in a dimer form (41). Our living cell homo-FRET approach allowed us to quantitatively measure the dynamic formation of the nanoclusters of plant surface molecules (39). Homo-FRET experiments revealed that flagellin triggered the nano-clustering of FLS2 from a less to more oligomerized, representing a transition from the unstimulated to active state during early immune responses.

Our data suggest a plausible model (Fig. 6) in which the *Xcc* QS molecule DSF alters sterols composition to modulate membrane microdomains and thereby, affects both the general endocytosis pathway as well as FLS2 receptor nano-clustering upon PAMP stimulation, and thus the ROS production (100). Sterols, together with sphingolipids, are major constituents of plasma membrane lipid raft/microdomains that are formed by diverse mechanisms of molecular assemblies of surface molecules (100–102). Distinct microdomains spatially separate from each other and their assembly and distributions are of significant importance for the regulation of diverse cellular processes, including the nano-clustering or CME of surface signaling factors (41, 103, 104). In agreement with this model and our data, sterol content has previously been found to directly or indirectly influence membrane organization and CME (41, 58, 104, 105). CME is well-known to coordinate PTI signaling in plants 106. However, as ligand-induced MAPK activation (~5-30 min post elicitation) is detected earlier than the appearance of endocytosis-mediated PRR-accumulation in endosomes (~30 min-1 h post elicitation) (18, 26) (Fig. 2A), endocytic invagination per se might not be a direct mediator for MAPK activation (24), which is also supported by the fact here that an attenuated-endocytosis by DSF did not alter flg22-mediated MAPK activation. Nevertheless, the impairment of endocytosis could still influence the internalization and recycling of the surface PRRs at different oligomerization states upon PAMP-elicitation (24, 107). Consistent with this notion, recent work also highlighted many consequences of the CME impairment in the *Arabidopsis* clathrin mutant *chc2*, such as FLS2 endocytosis, stomatal defense, callose deposition, as well as increased susceptibility to bacterial infection, while acute MAPK response remained intact (20).

**Fig. 6.**
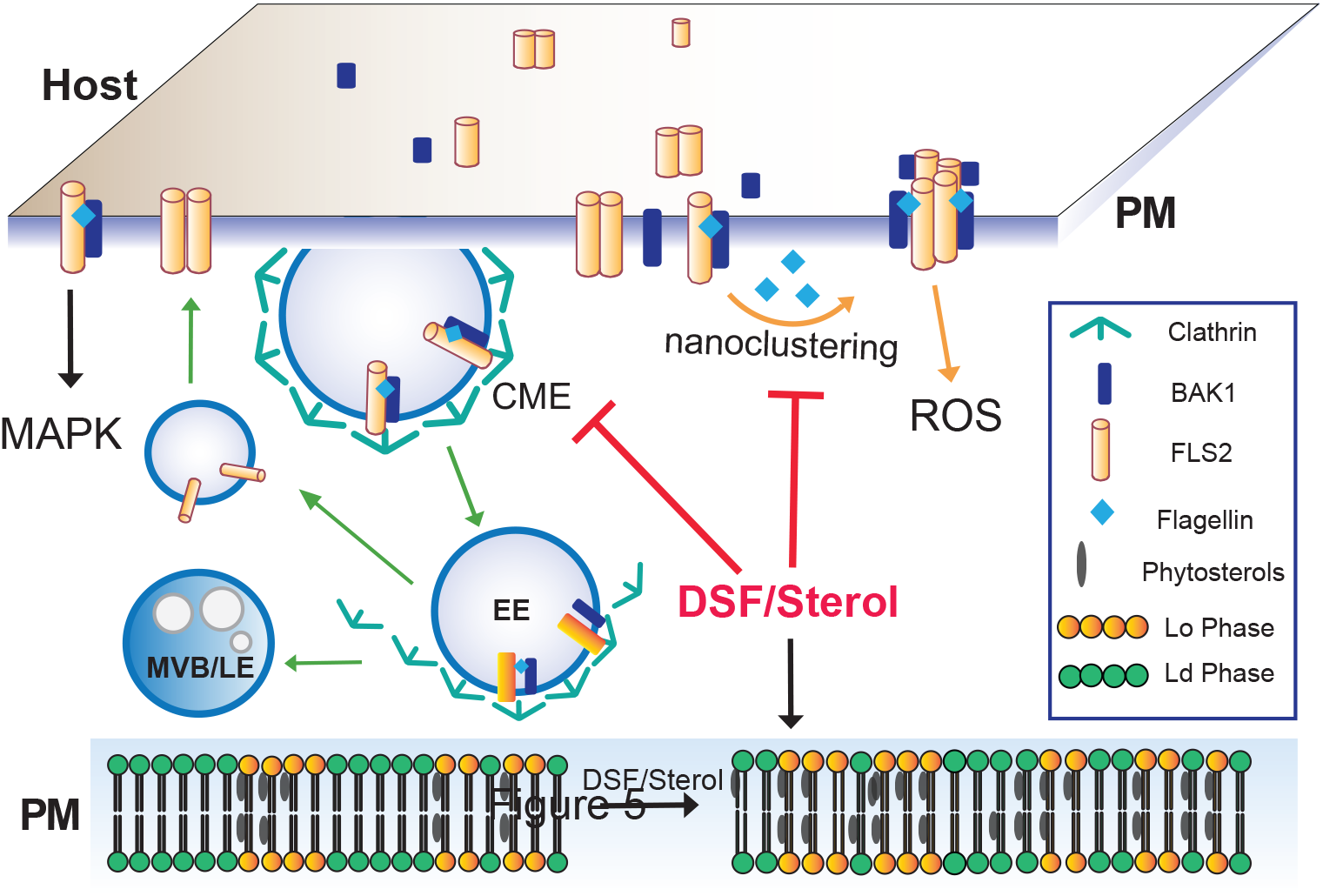
Proposed model for DSF-induced suppression of plant innate immunity by pertubation of PM lipids. Upon contact with bacteria/flagellin, FLS2 receptors form homodimers and heterodimers with interacting partners on the plasma membrane. Activated receptors then relay the signals to PM co-receptors and cytoplasmic kinases to induce a wide range of defense responses and also undergo endocytosis. However, in plants infected with DSF-producing bacteria, DSF could be metabolized into plant sterols or directly induce the production of phytosterols via unknown mechanisms. Sterol enrichment on the plant PM leads to an increase in lipid ordered phase - L_o_ (or PM microdomains) and a decrease in lipid disordered phase (L_d_ phase). Receptor nanoclustering were affected and as a result, they could not oligomerize and undergo endocytosis properly. Therefore, signalling cascades from plasma membrane that result in PTI responses were attenuated.

During bacterial propagation from early-to late-stages of infection, DSF attenuates the continuous enhancement of the host PTI responses in ROS production that could otherwise be stimulated by the increasing amount of PAMPs. Receptor activities in both lateral motility for nano-clustering or internalization for receptor recycling on the plasma membrane are compromised by DSF signalling. Our report emphasizes the role of QS molecules beyond intraspecies communication, which has important implications for plant disease control (3). It will also be of particular interest to investigate if the regulation of host metabolic pathways by different small molecules from pathogens is a ubiquitous phenomenon during host-pathogen interaction. In addition, since quorum-sensing molecules from human pathogens, such as *P. aeruginosa* have also been documented to disrupt host membrane microdomains (72), it is worth future studies to investigate whether signalling of plasma membrane receptors such as TLR5 (an ortholog of FLS2) is also affected by such molecule in a sterol-dependent manner.

## ACKNOWLEDGEMENTS

We are grateful to Kimberly Kline (Singapore Centre for Environmental Life Sciences Engineering, Singapore) for the critical reading of the manuscript. We thank Lay Yin Tang and Alma Turšić-Wunder for helping in RNA preparation. We thank the following researchers for sharing the *Arabidopsis* seeds: Takashi Ueda (National Institute for Basic Biology, Japan) for the FLS2-GFP *Arabidopsis*, Jianwei Pan (Lanzhou University, China) for PIN2-GFP *Arabidopsis*, Liwen Jiang (Chinese University of Hongkong) for the VHAa1-GFP line, and Thomas Ott (University of Freiburg) for the pREM1.2::YFP:REM1.2 line; Kathrin Schrick (Kansas State University), Jyan-Chyun Jang (Ohio State University) for the *Arabidopsis* sterol mutants; and Jinbo Shen (State Key Laboratory of Subtropical Siviculture, Zhejang A&F University, China) for the BOR1-GFP line. We also thank Yinyue Deng (South China Agricultural University, China) for sharing reagents, M. K. Mathew and Divya Rajagopal (National Centre for Biological Sciences-TIFR, India) for helping with *Arabidopsis* growth for the Homo-FRET experiments. SM is supported by a Margdarshi Fellowship (IA/M/15/1/502018) from the DBT-Wellcome Trust Alliance. This study was supported by NTU NIMBELS (NIM/01/2016) and NTU startup grant (M4081533) to Y.M. in Singapore.

## Contributions

T.M.T., Z.M., and Y.M. conceived and designed the study. T.M.T. and Z.M. performed most of the plant-based experiments. T.M.T., A.T., and F.T. performed lipidomics analysis and interpreted the data. Z.M. and S.N. performed homo-FRET based imaging, image analysis, and data interpretation. Y.C helped with the quantitative RT-PCR and data analysis. X.H. and J.W. performed the EM works. B.G. and J.L. performed the CoIP experiment. F.T., M.R., S.M., J.L., and L.Y. provided essential intellectual input and data interpretation. T.M.T. and Y.M. wrote the manuscript with feedback from all authors. Y.M. supervised the study.

---

**Fig. S1. DSF affected plant growth and flagellin receptor degradation (A)** Growth of *Arabidopsis* Col-0 seedlings seven days on 1/2 Murashige-Skoog agar supplemented with different concentrations of DSF (bar, 10 mm, n≥14). **(B)** Confocal micrographs of the FLS2 receptor signal on PM of FLS2-GFP seedlings after treatment with 10 μM flgII-28 (bar, 10 μm). Micrographs are maximum projections of 12 slices at every 1 μm z-distance. Bargraph on the right represent number of endosomes per image quantified from maximum-projected images (Error bars, SEM, n≥12). **(C)** Degradation of FLS2 receptor was delayed in DSF-treated plants. Leaf strips of 5-week old FLS2-GFP plants were floated for 24 h in water containing DMSO or 25 μM DSF, then elicited with 10 μM flg22 and snap-frozen in liquid nitrogen at indicated timepoints. Total protein extracts were probed with *α*-GFP antibody to detect FLS2. Staining of the membrane with Ponceau S served as loading control. The experiment was repeated three times with similar results **(D)** Western blot analysis of MAPK responses of Col-0 leaf strips in response to the flg22 peptide at different time-points, following 24 h treatment in 1/2 MS supplemented with DMSO or 25 μM DSF. Ponceau S staining of the WB membrane was shown as loading control. The experiment was repeated twice with similar result. **(E**) Relative transcript abundance of *FRK1* as measured by quantitative RT-PCR of *FRK1* expression. Data represent relative expression to control calculated using the ΔΔCt method. Columns represent mean relative gene expression level of at least six technical replicates from two biological replicates (Error bars, SEM). (**F**) Co-immunoprecipitation assay on *Arabidopsis* protoplast co-transfected with FLS2-HA and BAK1-FLAG constructs. Transfected protoplasts were treated with 1 μM DSF for 12 h and elicited with 1 μM flg22 for 10 min and CoIP was performed with anti-FLAG antibody. The experiment was repeated twice with similar result. P-values were determined by one-way ANOVA (^∗^ p < 0.05; ^∗∗^ p < 0.01; ^∗∗∗^ p < 0.001; ^∗∗∗∗^ p < 0.0001; ns, not significant).

**Fig. S2. Receptor endocytosis and clustering is affected by DSF**

**(A-C)** Kymographs and lifetime of clathrin light chain (CLC-GFP, n≥250 events), brassinosteroid receptor BRI1 (BRI1-GFP, n≥136 events) and Boron receptor BOR1 (BOR1-GFP, n≥198 events). Kymographs were derived from VA-TIRF images obtained from corresponding marker lines grown on ½ MS plates for 5-6 days, then treated with DMSO (control) or 25 μM DSF for 24 h in ½ MS. **(D)** Anisotropy measured for purified GFP protein solution at definitive TIRF (incident angle was set as 1880), and two HILO angles (1910 and 2000). Quantification data shows similar anisotropy values can be obtained in these three different incident angles (bar, 2 μm, n≥42 ROIs). **(E)** Anisotropy measured at HILO-TIRF gave the same result as definitive TIRF, in which samples could be illuminated evenly. Anisotropy of control and M*β*CD (10 mM in Glucose M1 buffer for 45 min at 37°C) treated cells were measured at definitive TIRF (1880) and HILO-TIRF (1950 and 2000) using stably transfected GFP-GPI construct in CHO cells. Measurements show the same results at definitive TIRF (1880) and HILO-TIRF (1950); and at HILO-TIRF (2000) similar values can be obtained if measurements were considered from evenly illuminated area, which is highlighted by dotted line (bar, 5 μm, n≥30 ROIs). P-values were determined by one-way ANOVA (^∗^ p < 0.05; ^∗∗^ p < 0.01; ^∗∗∗^ p < 0.001; ^∗∗∗∗^ p < 0.0001; ns, not significant).

**Fig. S3**. **DSF did not cause morphological changes in *Arabidopsis* endosomal compartments. (A-C)** Confocal micrographs of DMSO/DSF-pretreated *Arabidopsis* early endosome marker line (VHAa1-GFP), late endosome marker line (RabF2b-YFP), and Golgi marker line (Got1p-YFP) stained with FM4-64 dye with or without BFA treatment (bar, 10 μm). The relative size of enlarged multivesicular bodies (MVB) and BFA compartments were measured by Fiji. **(D)** Ultrastructural observation of *Arabidopsis* organelles in eight-day-old seedlings pre-treated with DMSO/DSF observed by Transmission Electron Microscopy (upper panel scale bar, 200 nm; lower panel scale bar, 500 nm). Annotations on the electron-micrographs are as followed: G, Golgi; TGN, trans-Golgi network; CW, cell wall; m, mitochondria; MVB, multi-vesicular body; C, Chloroplast. **(E)** Sterol biosynthetic pathways in *Arabidopsis thaliana*. Corresponding mutants were labelled in each step of the pathways, with dashed arrows indicating multiple biosynthetic steps. **(F)** Growth of sterol biosynthesis mutants *fk-X224 and smt1-1* and their corresponding wildtype ecotypes (Ler and Ws, respectively) on ½ MS agar supplemented with DMSO or 25 μM DSF. Plates were scanned after seven days after sowing the seeds (bar, 5mm). The experiment was repeated twice with similar result. P-values were determined by 2-tailed Student’s *t*-test (^∗^ p < 0.05; ^∗∗^ p < 0.01; ^∗∗∗^ p < 0.001; ^∗∗∗∗^ p < 0.0001; ns, not significant).

**Fig. S4. Sterol removal by M*β*CD restored phytosterol-induced inhibition of endocytosis. (A)** FM4-64 dye uptake by PIN2-GFP roots treated with M*β*CD in combination with ES9-17/BFA (n≥5; bars, 10 μm) **(B)** Life time of CLC-GFP in the presence of DSF, M*β*CD and a combination of both drugs. CLC-GFP seedlings were treated with DMSO/25 µM DSF for 24 h in ½ MS medium, then subjected to a brief treatment of 2 mM M*β*CD before being imaged using VA-TIRF. **(C-D)** Sterol mix similarly inhibits FM4-64 uptake in combination with the clathrin inhibitor ES9-17. Five-day old PIN2-GFP seedlings were transferred to ½ MS plates containing 50 µM sterol mix **(C)** or 50 µM sterol mix + 10 mM M*β*CD **(D)** for an additional day. Plants were subjected to drug treatments (50 µM ES9-17–1h, 50 µM BFA – 1h, or a combination of both drugs) before being stained with FM4-64 and imaged (n≥5, bars, 10 μm). P-values were determined by one-way ANOVA (^∗^ p < 0.05; ^∗∗^ p < 0.01; ^∗∗∗^ p < 0.001; ^∗∗∗∗^ p < 0.0001; ns, not significant).

**Fig. S5. Phytosterols mimic DSF-induced inhibition of receptor internalization, ROS production and primary root growth**

**(A)** Internalization of the FLS2 receptor was monitored in FLS2-GFP seedling cotyledons. Plants were pre-treated with 50 µM sterol mix or sterol mix + 2 mM M*β*CD for 24 h prior to elicitation with 10 μM of flg22 and imaged at indicated timepoints. (n≥12, bars, 10 μm). Boxplots represent the number of endosomes per image as quantified by segmentation and particle analysis using Fiji. **(B)** ROS burst in response to flg22 in Col-0 leaf strips pre-treated for 24 h with solvent control or 50 μM sterol mix, with or without 2 mM M*β*CD. **(C)** ROS production by Col-0 leaf strips pre-treated with DMSO/25 μM DSF for 24 h simultaneously with the sterol biosynthesis inhibitor Lovastatin (Lova, 1 μM). Leaf strips were then elicited with 1 μM flg22 and luminescence was monitored every minute for 1 h by a BioTek Cytation 5 plate-reader. The experiment were repeated twice with similar result. **(D)** Root growth of Col-0 plants on ½ MS plates supplemented with different concentrations of *β-*Sitosterol seven days after sowing (n≥22). **(E)** Dose-dependent reversion of *β-*Sitosterol-induced growth inhibition of *Arabidopsis* Col-0 by M*β*CD. Col-0 seeds were germinated on ½ MS agar supplemented with *β-*Sitosterol in combination with 2 mM or 10 mM of M*β*CD. After seven days, the plates were scanned and primary root lengths were measured using Fiji. Boxplot represents data from n≥20 seedlings. P-values were determined by one-way ANOVA (^∗^ p < 0.05; ^∗∗^ p < 0.01; ^∗∗∗^ p < 0.001; ^∗∗∗∗^ p < 0.0001; ns, not significant).

**Fig. S6. Osmotic stress relieved DSF-induced inhibition of endocytosis in *Arabidopsis*. (A)** Growth of Col-0 seedlings on 1/2 MS agar supplemented with DMSO or 25 μM DSF in combination with increasing concentrations of NaCl or Mannitol. Seedlings primary root length was measured after 7 days growing on agar plates (n≥23 seedlings). **(B)** Recovery of FLS2 internalization into endosomes by NaCl. Six-day old FLS2-GFP seedlings were transferred to liquid ½ MS supplemented with DMSO or 25 μM DSF for 24 h, then treated with 100 mM NaCl for 1h prior to flg22 elicitation and imaged at the indicated time points. Micrographs represent maximum projection of 12 slices taken every 1 μm (bar, 10 μm, n≥12 images). P-values were determined by one-way ANOVA (^∗^ p < 0.05; ^∗∗^ p < 0.01; ^∗∗∗^ p < 0.001; ^∗∗∗∗^ p < 0.0001; ns, not significant).

**Dataset S1.** Lipidomic profile of *Arabidopsis* Col-0 seedlings grown on ½ MS agar plates supplemented with DMSO, DSF, M*β*CD and DSF+ M*β*CD. Plants were grown vertically on ½ MS agar amended with appropriate chemicals in each treatment for seven days before tissues were collected for lipid extraction and analysis.

**Dataset S2.** Relative fold-change of lipidomic profile of *Arabidopsis* Col-0 seedlings grown on DSF, M*β*CD and DSF+ M*β*CD compared to DMSO control. P-values were determined by one-sample *t-test* to determine if sample means are different than 1 at the confidence interval of 90%.

**Table S1.** Key resource table

## CONTACT FOR REAGENT AND RESOURCE SHARING

Material used in this study can be found in the Key Resource Table (Table S1). Further information and requests for reagents and resources should be directed to and will be fulfilled by the Lead Contact, Yansong Miao

## EXPERIMENTAL MODEL AND SUBJECT DETAILS

### Plant growth

*Arabidopsis* [*Arabidopsis thaliana* (L.) Heynh.] plants were maintained in growth chamber at 22°C under long-day condition (16 h light/8 h dark). For *Arabidopsis* plants used in ROS assay and Western blot, plants were kept at short-day condition (8 h light/16 h dark) to facilitate vegetative growth. Col-0, RabF2b-YFP (Wave2Y), Got1p homolog-YFP (Wave 22Y) seeds were obtained from the Arabidopsis Resource Center (ABRC) stock centre (Ohio, U.S.A.). *Arabidopsis* FLS2-GFP, pREM1.2::YFP:REM1.2, CLC-GFP, BRI1-GFP and BOR1-GFP marker lines were described previously (18, 57; 108, 109,110). Sterol mutants *fkX224*, *smt1-1* and their WT ecotypes were described previously (111, 112). For cell biology imaging, *Arabidopsis* seeds were surface sterilised with 10% bleach and 70% EtOH, washed three times with sterilised water and vernalized at 4°C for 2 days. Seedlings were then sown on ½ MS agar and grown at 22°C for 4-5 days under long-day condition (16 h light/8 h dark) before treatments and imaging. For growth assay to evaluate primary root growth, we germinated surface-sterilised seeds directly on ½ MS agar containing indicated drugs. Plants were grown vertically at 22°C for seven days before images of the plates were taken using a flatbed scanner and primary root length was determined by Fiji software. Unless specified, all growth assays were repeated at least twice with approximately 20-30 seedlings for each treatment. For growth assay of sterol biosynthesis mutants on DMSO- and DSF-supplemented medium, growth inhibition rate (%) were defined as the percent difference between the root length of DSF-treated plants and the Mean root length of DMSO-treated plants.

### Bacterial strains

*Pseudomonas syringae* pv. *tomato* DC3000 (*Pst*) (113) were cultured at 28 °C on NYG medium.

## METHODS

### Chemicals and treatment conditions

DSF (*cis*-11-methyl-2-dodecenonic) was purchased from Merck and dissolved in DMSO to obtain 100 mM stock. The bacterial flagellin peptides flg22 and flgII-28 were chemically synthesised (GL Biochem Ltd., China) with 95% purity and dissolved in water to obtain 5 mM stocks. For pharmacological studies, Brefeldin A (BFA), Wortmannin, Lovastatin, and ES9-17 were dissolved in DMSO as 50 mM stocks. Unless specified, seedlings were incubated with these drugs at the working concentration of 50 μM in ½ MS medium for 1 h at room temperature. The sterol-depleting reagent M*β*CD was dissolved in deionised water at 200 mM and filter-sterilized. For short-term M*β*CD treatment, *Arabidopsis* seedlings were treated with 2 mM M*β*CD in liquid ½ MS for 30 min at room temperature. Long-term growth assay with M*β*CD was performed by germinating axenic *Arabidopsis* seeds directly on ½ MS agar containing 2mM M*β*CD.

### qRT-PCR

*Arabidopsis* Col-0 seeds were surface-sterilized and vernalized as mentioned above. Five-day old seedlings were transferred to new ½ MS plates containing either DMSO or 25 μM DSF for three additional days. At day 8, the seedlings were subjected to a brief treatment with either 150 nM of flg22 or water control and plant tissue was frozen in liquid nitrogen and kept at −80°C. RNA extraction was performed using RNeasy Plant Minikit (Qiagen) according to manufacturer’s instruction. Each treatment includes 2-3 biological replicates.

DNase treatment was performed twice using Turbo DNase (Ambion) and RNAClean XP kit (Agencourt). The quality of RNA samples was evaluated using the Qubit Fluorometer (Thermo Fisher). cDNA was synthesized by SuperScript III First-Strand Synthesis system (Invitrogen), using oligo-d(T) primer. Quantitative RT-PCR was performed in triplicate for each of the biological replicates, using Kapa SYBR FAST qPCR Master mix with primers specific for *FRK1* (13), and *EF-1α* gene used as an internal control (114). Data were collected by Applied Biosystem StepOnePlus RealTime PCR system (Applied Biosystem) and analyzed using the manufacturer’s software.

### Microscopy and data analysis

#### Confocal microscopy

Confocal microscopy was performed on a Zeiss ELYRA PS.1 + LSM780 system equipped with a 63X Plan-Apochromat oil-objective (NA=1.4). GFP/YFP and FM4-64 were excited with a 488-nm and 514-nm laser, respectively. Emission was collected at 493-594 nm for GFP/YFP and 612-758 nm for FM4-64, respectively. In addition, chloroplasts autofluorescence in FLS2-GFP cotyledons was also captured from 630-730 nm.

For imaging of FLS2-containing endosomes, image acquisition was performed as described previously (115). Drug treatments of FLS2-GFP seedlings were performed in liquid ½ MS as followed: DSF: 25 μM-24 h; Sterol mix (Sitosterol:Stigmasterol:Campesterol = 8:1:1): 50 μM-24 h; M*β*CD: 2 mM - 30 min (alone or in combination with 25 μM DSF-24 h); 10 mM – 24 h (alone or in combination with 50 μM sterol mix, 24 h). After drug treatments, FLS2-GFP seedlings were further elicited with 10 μM flg22 and imaged at 30’, 60’ 120’ post-elicitation (water was used as mock control). We also included at treatment of 10 μM flgII-28 as negative control since flgII-28 peptide is not recognized by *Arabidopsis* FLS2 receptor. At indicated timepoints, z-stack images of FLS2-GFP seedling cotyledons were taken at 12 slices of 1 μm z-distance. Endosome signals from maximum projection images were segmented to remove plasma membrane signal using Fiji “Trainable Weka Segmentation” plug-in. Segmented images were analyzed further by “Particle Analysis” tool in Fiji (116) to obtain the number of endosomes per image. Experiments were repeated twice, with 12 images taken from three seedlings for each replicate.

To monitor FM4-64 uptakes in PIN2-GFP roots, seeds were sown on ½ MS agar and grown vertically. Five-day old PIN2-GFP seedlings were then transferred to a new plate containing either DMSO (control), 25 μM DSF or 50 μM sterol mix (Sitosterol:Stigmasterol:Campesterol = 8:1:1) for an additional day. After 24 h of growth on DMSO/DSF/Sterol-supplemented media, the following drug treatments were performed in liquid ½ MS at room temperature: M*β*CD: 2 mM - 30 min (alone or in combination with 25 μM DSF-24 h treatment); M*β*CD: 10 mM – 24 h (alone or in combination with 50 μM sterol mix-24 h treatment); NaCl: 100 mM - 1h; Mannitol 100mM - 1h; ES9-17: 50 μM-1h; BFA: 50 μM-1h. After treatments, PIN2-GFP seedlings were incubated with 2 μM FM4-64 for 5 min at room temperature, washed twice with fresh ½ MS medium and roots were imaged immediately using an ELYRA PS.1 + LSM780 system equipped with a 63X Plan-Apochromat oil-objective (NA=1.4).

#### VA-TIRFM and lifetime analysis

Variable-angled Total Internal Reflection Fluorescence Microscopy was performed on a Zeiss ELYRA PS.1 + LSM780 system equipped with a 100X Plan-Apochromat oil-objective (NA=1.46) to monitor the dynamics of PM markers. Time-lapse images of FLS2-GFP foci on plasma membrane was taken at up to 120 frames with 1000-ms exposure at 2% laser power. Time-lapse movies of other PM marker lines were acquired as followed: CLC-GFP: 90 cycles, 1000-ms exposure, 1000-ms interval, BRI1-GFP and BOR1-GFP: 90 cycles, 500-ms exposure, 500-ms interval. Acquired VA-TIRFM images were deconvolved using Huygens Essential (Scientific Volume Imaging) using theoretical PSFs. Particle lifetime was measured from kymographs in Fiji. Lifetime of at least 100 individual foci from three different seedlings were presented. The clustering of FLS2-GFP foci on plant plasma membrane, indicated by spatial clustering index (SCI) was measured as described previously (117). Briefly, SCI was calculated based on the ratio of top 5% pixels with the highest intensity and 5% of pixels with the lowest intensity along 10-μM line ROIs selected from the first frame of each VA-TIRFM images. Data presented in boxplots were from at least 60 individual ROIs.

#### 2D-Structure Illumination Microscopy

2D-SIM (Structured Illumination Microscopy) image for pREM1.2::YFP:REM1.2 signals was conducted using a Zeiss ELYRA PS.1 + LSM780 system with a Zeiss Alpha Plan Apochromat 100X (NA=1.46) oil objective. A 488 nm laser light source was used to excite the YFP signal and the emission was collected in the range of 495-575 nm by a sCMOS camera (PCO) with a pixel size as 6.5 μm x 6.5 μm, the exposure time was set as 300 ms. All the 2D-SIM images contained five rotations and five phases of the grated pattern (42 μm) for each image layer. SIM image reconstruction was conducted by setting the Super-Resolution (SR) frequency weighting as 0.5 and the noise filter as −4.

Particle tracking was performed in Imaris (Bitplane). Imaris Spots function was used to automatically quantify the REM1.2 particle signals. To trace all the spots, the estimated spot diameter was set to 0.22 μm according to the preliminary particle analysis before processing the whole image, the threshold was set to automatic region threshold. After tracing all the particles in each image, the total intensity of each particle were analyzed and output automatically. For each treatment, data of 500 particles were randomly selected from more than 10 images (more than 2000 particles in total for each treatment) for further statistical analysis.

#### FLS2-GFP steady-state anisotropy measurement

Steady state anisotropy were measured on FLS2-GFP expressed live cells in cytoledon using NikonTE2000 TIRF microscope with dual camera as described previously (38). Briefly, with respect to plane of polarization of 488 nm excitation light (~ 5.2 mW power), emission intensity was collected by two Evolve™ 512 EMCCD camera at parallel (I_pa_) and perpendicular (I_pe_) direction. Plasma membrane of the plant was illuminated at HILO using variable TIRF angle module until the best SNR was achieved using a 100X (NA=1.45) objective with image plane pixel size of 106 nm and 1 s exposure time. Anisotropy measured at HILO (1910, 1950 angles) showed same results as definitive TIRF, using GFP solution and stable transfected GFP-GPI construct in CHO cells (Fig. S2D-E). Anisotropy values measure at HILO (2000 angle) even showed similar results if ROI were taken from evenly illuminated area of the cells. However, we have restricted our all measurements at HILO angle <1950 to nullify any bias due to uneven illumination. G-Factor of the instrument was calculated with freely diffused FITC solution by calculating ratio of I_pa_ vs I_pe_ emission intensities (Ghosh et al., 2012). Then anisotropy of the FLS2-GFP was calculated by G-factor corrected perpendicular images using the following equation:

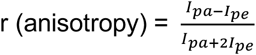

Alignment of images of I_pa_ and I_pe_ and G-factor corrections were done using algorithm written in Matlab (MathWorks). FLS2-GFP anisotropy in different treatment conditions was compared using data from more than 150 ROIs (20×20 pixel) from at least eight images taken from at least three seedlings. The experiment involved two biological replicates.

### Transmission electron microscopy

*Arabidopsis* Col-0 seeds were surface-sterilized and vernalized at 4°C for two days, then grown on ½ MS plates at 22°C (16 h light/8 h dark). After five days, the seedlings were transferred to a new ½ MS plates containing either 25 μM DSF or DMSO (solvent control) for an additional three days. Eight-day-old seedlings were severed into root, hypocotyl and cotyledon sections. Plant tissues were fixed in primary fixative (1 ml of 2.5% Glutaraldehyde prepared in 25 mM Sodium Cacodylate buffer pH 7.2) overnight at 4°C. Plant tissues were then washed four times in 25 mM sodium cacodylate buffer, 10 min each time. Then, plant tissues were submerged in 1 ml of secondary fixative (2% Osmium tetraoxide, 0.5% Potassium ferricyanide in water) for 2 h at room temperature.

The samples were washed four times with 1 ml of deionized water, each time for 10 min, then transferred to 2% uranyl acetate solution and incubated at room temperature for 4 h. Plant samples were washed four times with deionized water, each time for 10 min until the solution became clear. Sample dehydration was done at room temperature with a series of increasing acetone solution as followed: 30% - 20 min; 50% - 20 min; 75% - 20 min; 100% - 20 min (2X).

Tissue embedding was performed using Spurr Low-viscosity Embedding Kit (Sigma-Aldrich). Samples were incubated in increasing Spurr solutions, each time for 1 h as followed: 25%, 50%, 75% and 100%. After incubating with 100% Spurr, the samples were kept in fresh 100% Spurr solution overnight at 4°C. The next day, fresh 100% Spurr was added to the plant samples one more time. Plant tissues were then transferred into BEEM embedding capsules (Electron Microscopy Sciences) with the position of the samples at the bottom of the capsules, then incubated for 4 h at RT, followed by another 2 days at 60°C. Sectioning was performed using a Cryo Leica EM UC7 diatome (Leica, Germany) with sample thickness set to 70 nm.

Post-staining ultrathin sections were done by incubating the sections with uranyl acetate (in methanol) for 1 min, then washed with double deionised water. The sections were then incubated with Reynold’s lead citrate solution for 1 min and subsequently washed with deionised water, then dried in a fume hood. The plant tissue sections were imaged using a Hitachi Transmission Electron Microscope HT7700 (Hitachi, Japan) with the acceleration voltage of 80 kV.

### Lipidomic profiling

#### Lipid extraction

Surface-sterilized Col-0 seeds were directly germinated and grown vertically on ½ MS agar supplemented with DMSO, 25 *μ*M DSF, 2 mM MβCD or DSF+MβCD for 7 days at 22°C, 16 h light/ 8 h dark. After one week, 30 seedlings were pooled together as one biological replicate and total lipid was extracted as previously described (118). Prior to lipid extraction, 500 pmol each of PC 28:0, PE 28:0, PS 28:0, PG 28:0, PA 28:0, LPC 20:0, LPE 14:0, LPA 17:0, SM 12:0, Cer 17:0, CerOH 12:0, HexCer 8:0, TG 48:0 d5, GB3 23:0, and cholesterol d6 were added to each sample as internal standards. The experiment involved six biological replicates for each treatment.

#### LC-MS analysis

Dried lipid extracts were resuspended in 500 μL Butanol-Methanol (1+1, v/v) and stored at −20 °C until LC-MS analysis. Phospholipids, sphingolipids, and sterol lipids were identified using an Agilent 1290 UHPLC system coupled to an Agilent 6495 triple quadrupole mass spectrometer. Chromatographic separation was performed on an Agilent Eclipse Plus C18 reversed phase column (50 x 2.1mm, particle size 1.8 μm) using gradient elution of solvent A (water/acetonitrile 60/40, v/v, 10 mM ammonium formate) and solvent B (isopropanol/acetonitrile 90/10, v/v, 10 mM ammonium formate). 20% solvent B was increased to 60 % B over 2 min, and to 100 % B over 5 minutes. The column was flushed with 100 % B for 2 minutes, and then reequilibrated with 20 % B for a further 1.8 minutes. The flow rate was 0.4 mL/min, column temperature was 50 °C, and injection volume was 1 μL. The JetStream ion source was operated at the following conditions: Gas temperature, 200 °C; gas flow, 12 L/min; nebulizer gas pressure, 25 psi; sheath gas heater temperature, 400 °C; sheath gas flow, 12 arbitrary units; capillary voltage, 3500 V for positive ion mode, and 3000 V for negative ion mode; Vcharging, 500 for positive ion mode, and 1500 for negative ion mode; positive high pressure RF, 200; negative high pressure RF, 90; positive low pressure RF, 100; negative low pressure RF, 60; fragmentor voltage, 380 V; cell accelerator voltage, 5 V; dwell time, 1 ms.

The injection order of 24 samples (six replicates of four treatment groups) was randomised, and bracketed by injections of pooled quality control and extracted blank samples to monitor system stability and carryover. The whole sample set was analysed twice consecutively with different SRM settings. One injection was for phospholipids and sterols using SRM transitions from Vu *et al.* (118), and the second method detected plant sphingolipids with SRM transitions adapted from Markham et al. (119).

#### Data processing

Lipids were analysed based on characteristic SRM transitions and retention time using Agilent MassHunter Quantitative Analysis for QQQ (v. B.07.00). Further processing, including normalisation to suitable internal standards was performed in Microsoft Excel. High-resolution LC-MS data was processed using XCalibur QuanBrowser 3.0.63 (Thermo Fisher).

### Immunoblot analysis

Three leaves discs (0.7 cm in diameter) from 5 to 6-week old *Arabidopsis* plants were cut into strips and floated for 24 h in sterile water in 24-well plates under constant light to reduce stress due to wounding. DMSO or DSF were added into the wells 24 h before elicitation with 10 μM flg22. Leaf tissue was washed with sterile water to remove the elicitor and flash-frozen in liquid nitrogen at different timepoints. Total protein extraction was performed as described previously (85).

To detect the flagellin receptor, we used α-GFP antibody (1:200; Torrey Pines BioLabs, NJ, U.S.A) to probe for the FLS2-GFP in FLS2-GFP transgenic plants. For monitoring p-MAPK response, we used α-p-MAPK antibody (1:3000; Cell Signalling Technology, MA, U.S.A) to detect phosphorylated MAPKs in Col-0 leaf tissue after flg22 peptide elicitation. Western blot membranes were stained with Ponceau S as internal loading control.

### Measurement of callose deposition

Measurement of plant callose deposition in response to bacterial flagellin peptide was performed on two-week old Col-0 seedlings germinated on ½ MS medium. Whole seedlings were individually placed in 6-well plates with 4 ml of liquid ½ MS supplemented with DMSO or 25 μM DSF for 24 h at 22°C, then elicited for 24 h with 1 μM of flg22. Challenged plants were destained with acetic acid:ethanol (1:3) overnight, then washed in 150 mM K_2_HPO_4_ for 30 min. Destained leaves were incubated in 0.01% aniline blue (dissolved in 150 mM K_2_HPO_4_) in the dark for 2 h, then mounted on microscope slides in 50% glycerol. Callose deposition were observed on a Zeiss ELYRA PS.1 + LSM780 system using a DAPI filter (excitation:emission, 370:509 nm) with a 10X objective. The experiment was repeated twice, each involved 8 individual plants and at least 6 images were taken from each plant.

### Stomatal closure assay

Five-week-old Col-0 plants were used to determine the stomatal closure response in response to the flg22 peptide. Briefly, two-three leaves/plant were marked with a permanent marker, then infiltrated with DMSO or 25 μM DSF (diluted in sterilised 10 mM MgCl_2_). The plants were returned to the growth chamber for an additional 24 h. The next day, infiltrated leaves were excised from the plants and the abaxial epidermis was peeled off and transferred to a petri disc containing sterile water using a natural-hair brush (size 8). 10 μM flg22 was added to the samples for 30 min. The epidermal peels were stained with 20 μM propidium iodine for 5 min and rinsed briefly with sterile water, then mounted on microscope slides without coverslips and examined under a wide-field fluorescence microscope (Leica) equipped with RFP filter using a 10X objective lense. The experiments involved measurements of ≥130 stomata from at least 10 images taken from five individual plants.

### ROS production

Measurement of ROS burst in *Arabidopsis* apoplast elicited by flg22 was performed as described previously (24). Briefly, each leaf disc (0.7 cm in diameter) from 5 to 6-week old *Arabidopsis* plants was cut into 5 strips, floated on sterile water for 24 h in white 96-well plates under constant light at room temperature to reduce stress from wounding. Drugs were added into the wells prior to elicitation with 1 μM flg22 as followed: 25 μM DSF/DMSO control, 24 h; 2 mM M*β*CD, 30 min; 50 μM sterol mix, 24 h; 1 μM Lovastatin, 24 h. Each ROS experiment was performed in duplicate and experiments showing on the same graph were performed in the same plate.

### Plant infection assay

Two-week old *Arabidopsis* plants grown on ½ MS agar was used for flood-inoculation with *Pseudomonas syringae* pv. *tomato* DC3000 (*Pst* DC3000). Using *Pst* DC3000, a bacterial pathogen with a different quorum sensing system other than DSF, allowed us to dissect the effect of DSF on plant immunity independent of effector proteins activation and other bacterial traits regulated by quorum sensing. Briefly, plants were flooded with 40 ml of 10 mM MgCl_2_ solution containing 25 μM DSF or DMSO (control). After 1 day, the liquid was decanted and another 40 ml of MgCl_2_ containing either 1 μM of flg22 or water was poured into each plate. On the next day, the seedlings were inoculated with 5×10^6^ CFU/ml of *Pst* DC3000 prepared in 10 mM MgCl_2_. Bacterial population was determined 4 days post-inoculation (DPI) by dilution plating as described previously (120). The experiment was repeated twice with 9-12 technical replicates/treatment.

### Co-immunoprecipitation

The constructs for transient expression of BAK1 and FLS2 have been described earlier (93). Protoplast-based co-IP assays were conducted as previously described (121). Briefly, 1 ml of transfected protoplasts were incubated for 12 h for protein expression in the presence of 1 *μ*M DSF (Sigma, USA), and were treated with or without 1 *μ*M flg22 (MoonBiochem, China) for 10 min before harvest. Briefly, cells were lysed in 500 *μ*l IP buffer [10 mM HEPES, pH 7.5, 100 mM NaCl, 1 mM EDTA, 10% glycerol, 1% Triton-X100, and 1× protease inhibitor cocktail (Roche, USA)], and 30 *μ*l of protein lysate was kept as the input fraction. The remaining 470 *μ*l lysate was incubated with 10 *μ*l anti-FLAG M2 affinity gel (Sigma, USA) at 4°C for 3 h. After being washed five times with ice-cold IP buffer and once with 50 mM Tris-HCl (pH 7.5), the bound proteins were solubilized by boiling the beads in 60 *μ*l 2×SDS-PAGE loading buffer, and were detected by immunoblotting using anti-FLAG (Sigma, USA) or anti-HA antibody (Roche, USA). The experiment was repeated twice.

## QUANTIFICATION AND STATISTICAL ANALYSIS

Statistical analyses and data visualisation were performed using GraphPad Prism 7.0 (GraphPad, CA, U.S.A.). Mean comparisons were performed using one-way ANOVA (^∗^ p < 0.05; ^∗∗^ p < 0.01; ^∗∗∗^ p < 0.001; ^∗∗∗∗^ p < 0.0001; ns = not significant). Unless specified, data were presented as box-and-whiskers plots with bars indicated the median, 25% and 75% quartiles.

