## Supplemental figures and table for "The bacterial quorum sensing signal DSF hijacks *Arabidopsis thaliana* sterol biosynthesis to suppress plant innate immunity"

Figure S1

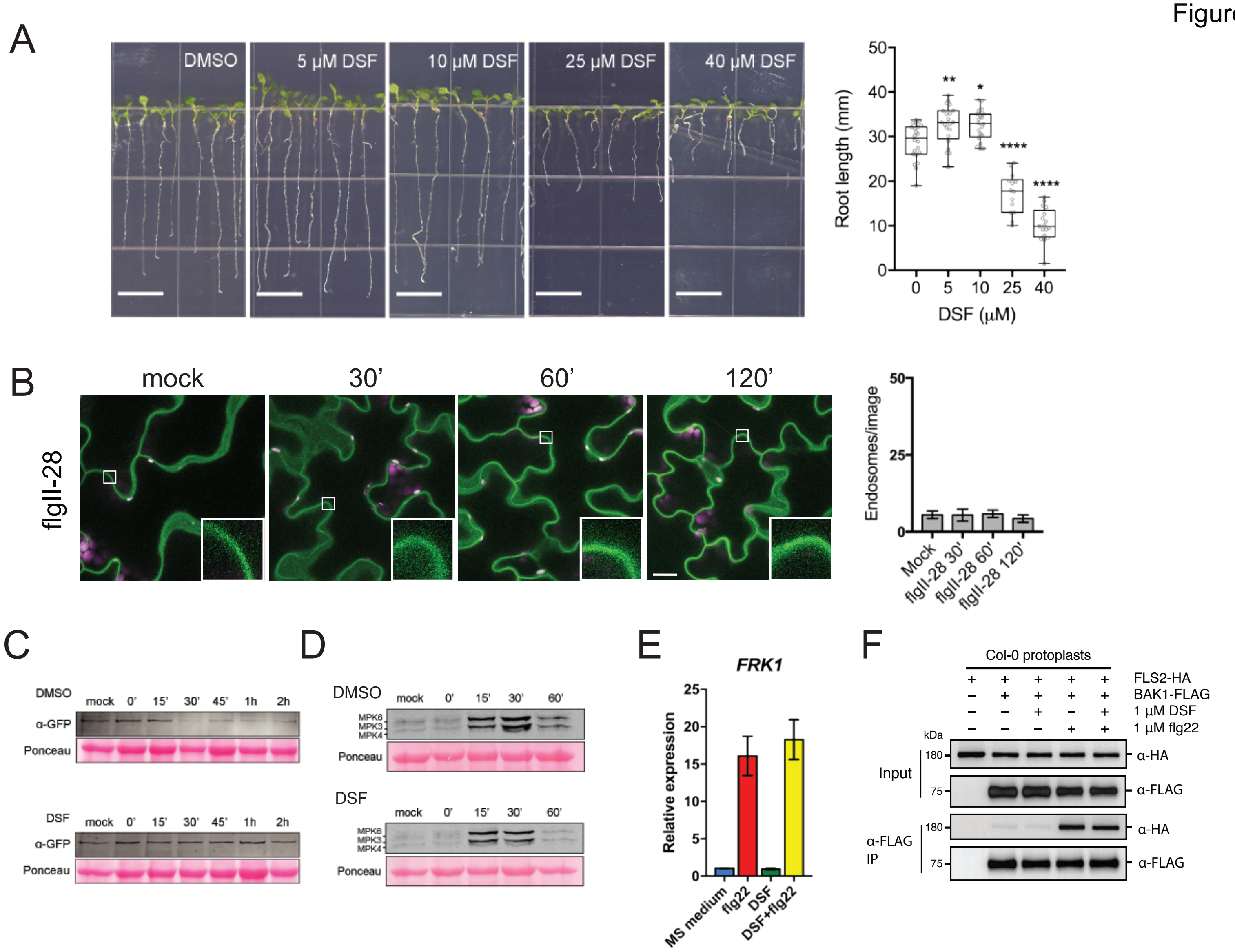

A
DMSO
5 µM DSF
10 µM DSF
25 µM DSF
40 µM DSF
Root length (mm)
DSF (µM)
B
mock
30'
60'
120'
flgII-28
Endosomes/image
Mock
flgII-28 30'
flgII-28 60'
flgII-28 120'
C
DMSO
DSF
mock 0' 15' 30' 45' 1h 2h
α-GFP
Ponceau
D
DMSO
DSF
mock 0' 15' 30' 60'
MPK6
MPK3
MPK4
Ponceau
E
FRK1
Relative expression
MS medium
flg22
DSF
DSF+flg22
F
Col-0 protoplasts
FLS2-HA
BAK1-FLAG
1 µM DSF
1 µM flg22
kDa
Input
α-FLAG IP
α-HA
α-FLAG

Figure S2

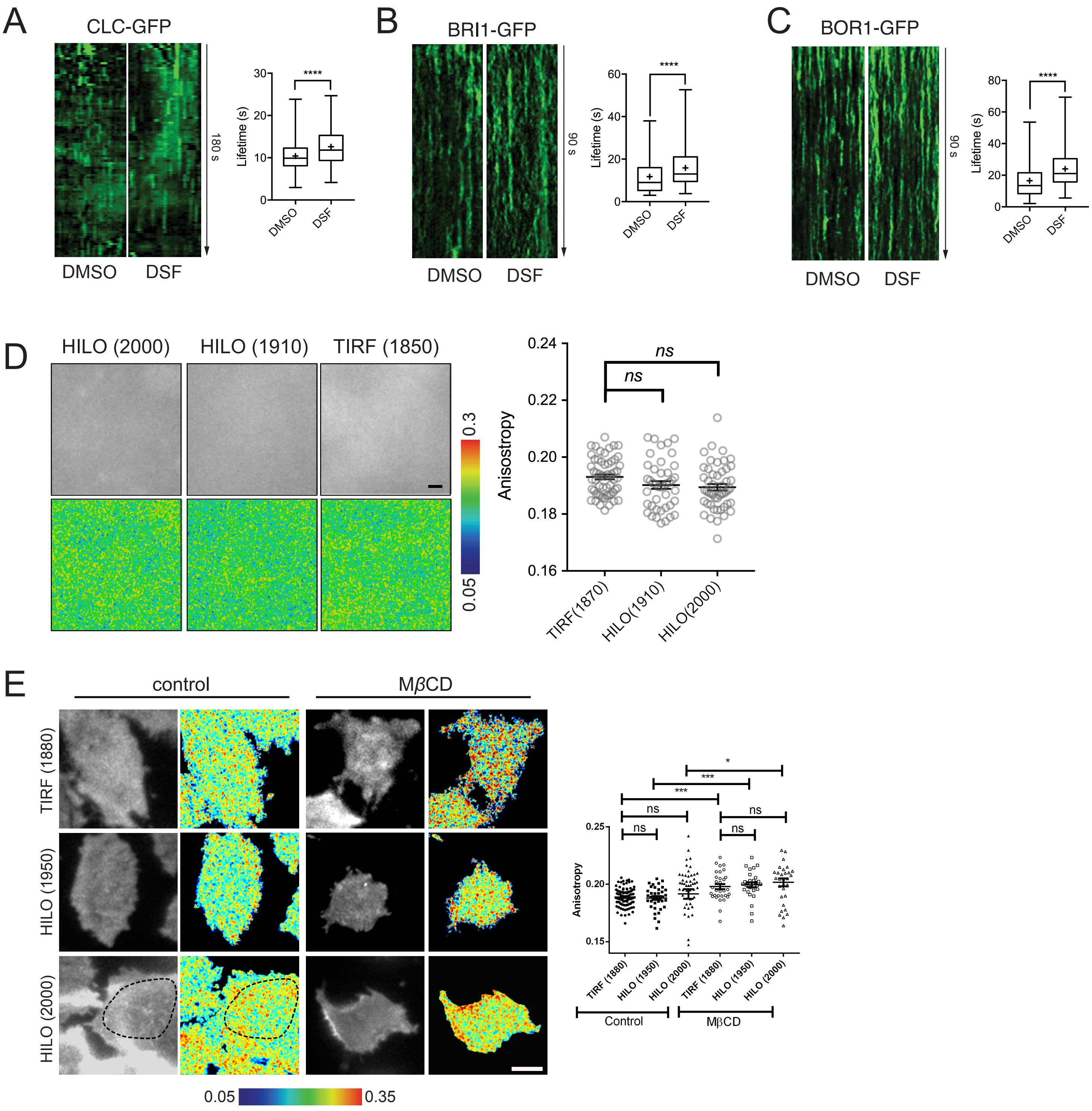

A
CLC-GFP
DMSO
DSF
180 s
Lifetime (s)
****
B
BRI1-GFP
90 s
C
BOR1-GFP
D
HILO (2000)
HILO (1910)
TIRF (1850)
0.3
0.05
Anisostropy
ns
TIRF(1870)
HILO(1910)
HILO(2000)
E
control
MβCD
TIRF (1880)
HILO (1950)
HILO (2000)
0.05
0.35
Anisotropy
ns
***
*
Control
MβCD

Figure S3

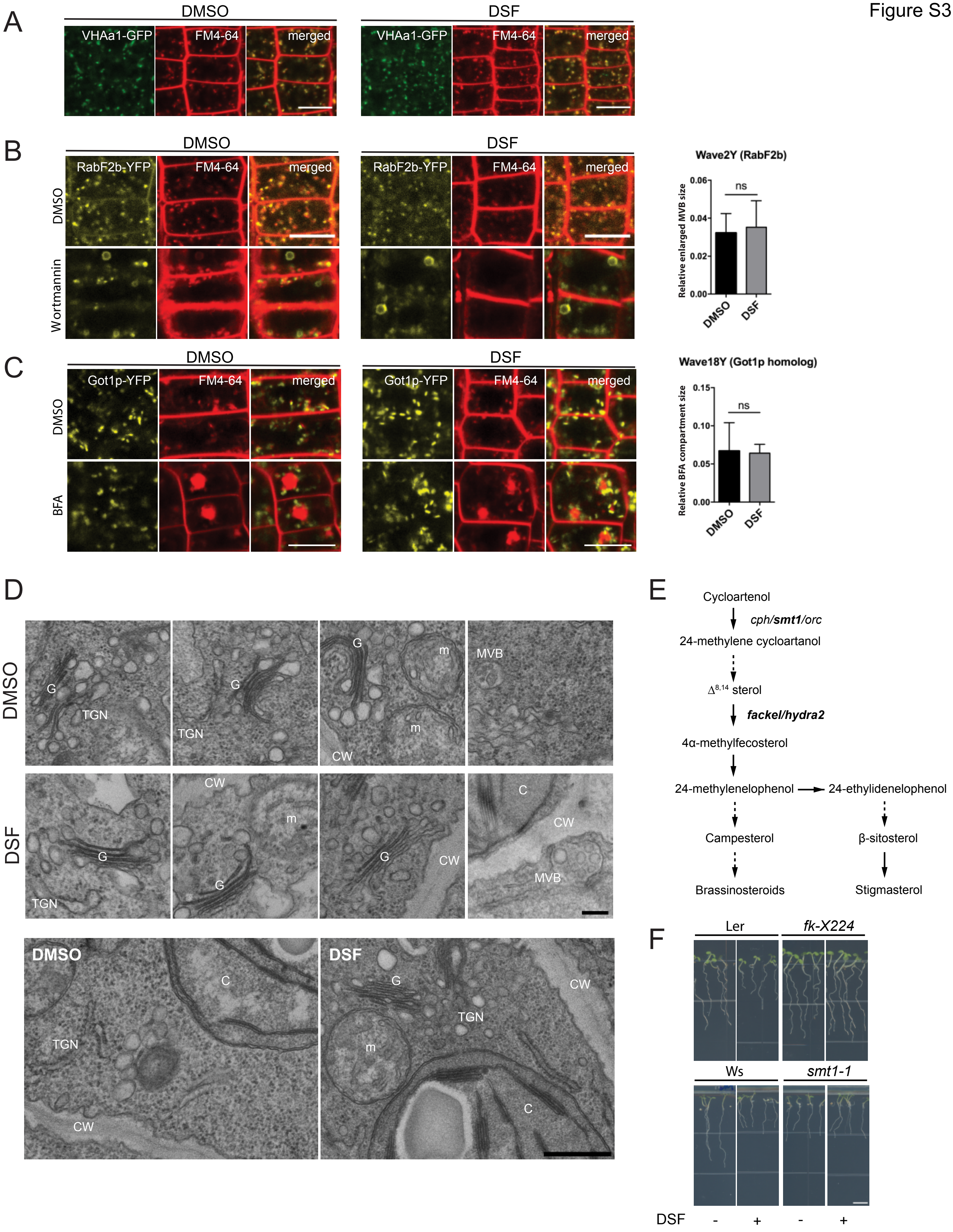
A
DMSO
DSF
VHAa1-GFP
FM4-64
merged
B
RabF2b-YFP
Wortmannin
Wave2Y (RabF2b)
Relative enlarged MVB size
0.06
0.04
0.02
0.00
ns
C
Got1p-YFP
BFA
Wave18Y (Got1p homolog)
Relative BFA compartment size
0.15
0.10
0.05
0.00
D
G
TGN
m
CW
MVB
C
E
Cycloartenol
cph/smt1/orc
24-methylene cycloartanol
Δ8,14 sterol
fackel/hydra2
4α-methylfecosterol
24-methylenelophenol
24-ethylidenelophenol
Campesterol
β-sitosterol
Brassinosteroids
Stigmasterol
F
Ler
fk-X224
Ws
smt1-1
DSF
-
+
-
+

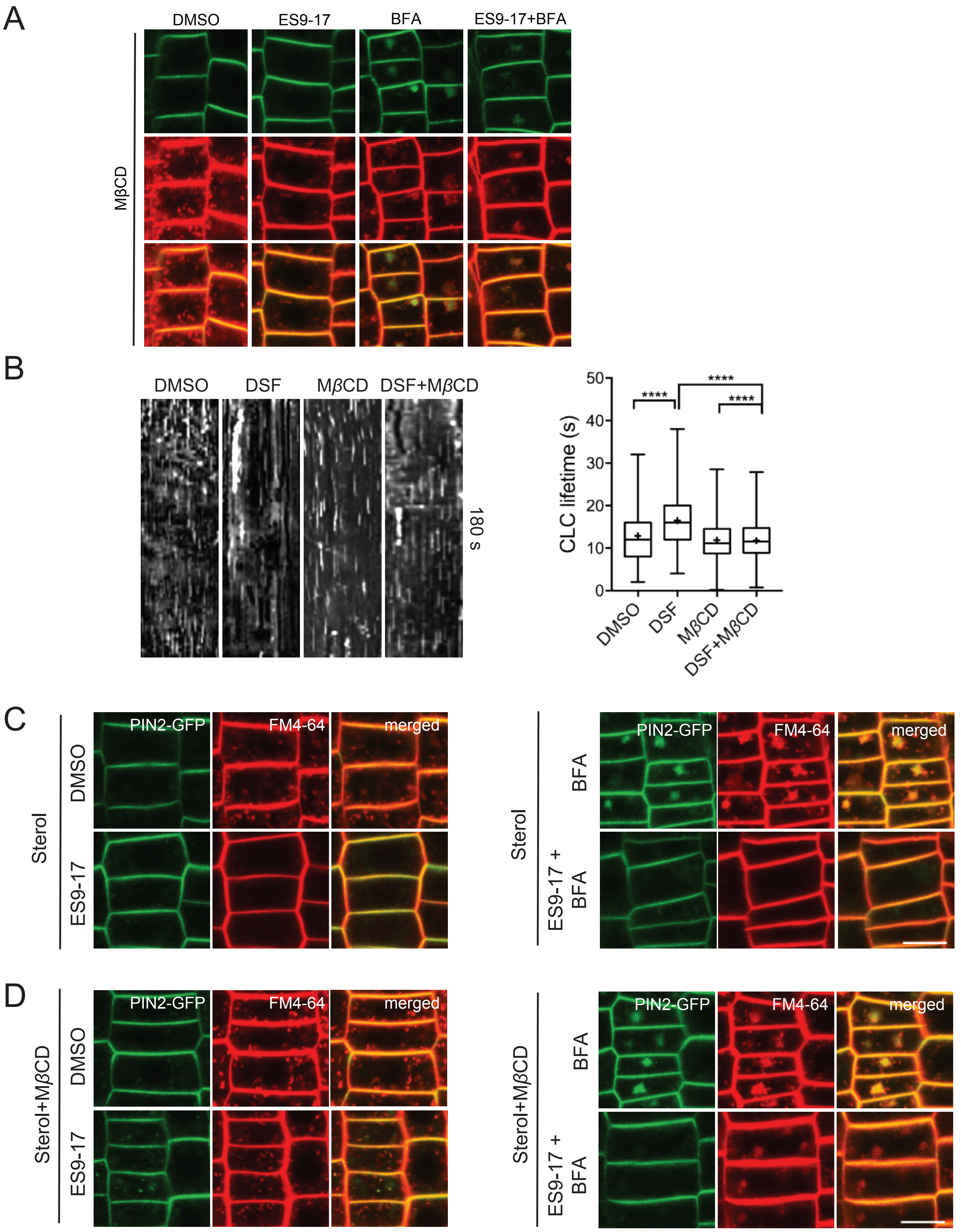
A
DMSO
ES9-17
BFA
ES9-17+BFA
MβCD
B
DMSO
DSF
MβCD
DSF+MβCD
180 s
CLC lifetime (s)
0
10
20
30
40
50
****
****
****
DMSO
DSF
MβCD
DSF+MβCD
C
PIN2-GFP
FM4-64
merged
Sterol
DMSO
ES9-17
BFA
ES9-17 + BFA
D
Sterol+MβCD
PIN2-GFP
FM4-64
merged
DMSO
ES9-17
BFA
ES9-17 + BFA

Figure S5

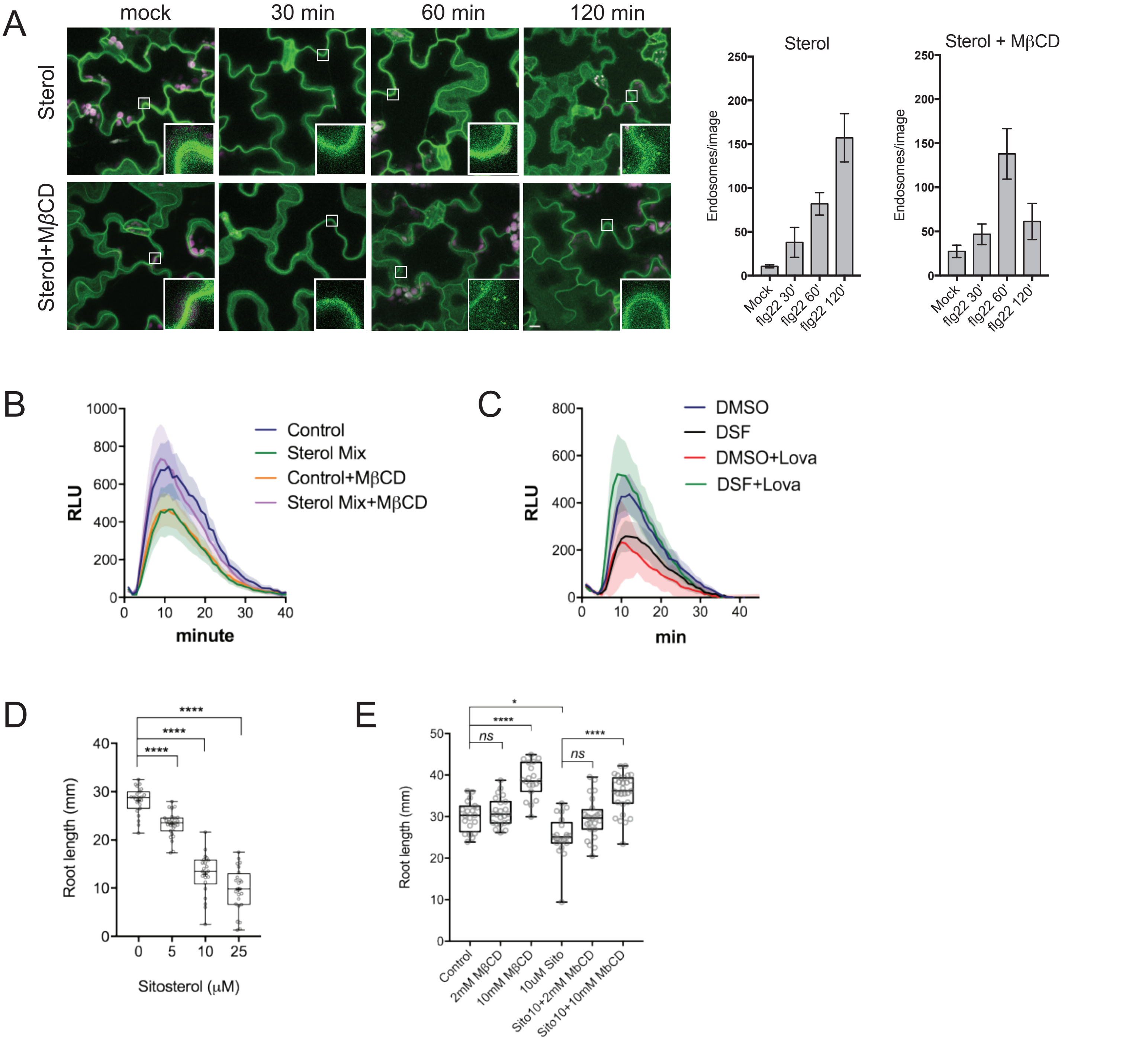

A
mock
30 min
60 min
120 min
Sterol
Sterol+MβCD
Sterol
Sterol + MβCD
Endosomes/image
Mock
flg22 30'
flg22 60'
flg22 120'
B
RLU
minute
Control
Sterol Mix
Control+MβCD
Sterol Mix+MβCD
C
RLU
min
DMSO
DSF
DMSO+Lova
DSF+Lova
D
Root length (mm)
Sitosterol (μM)
****
E
Root length (mm)
ns
****
*
Control
2mM MβCD
10mM MβCD
10uM Sito
Sito10+2mM MbCD
Sito10+10mM MbCD

Figure S6

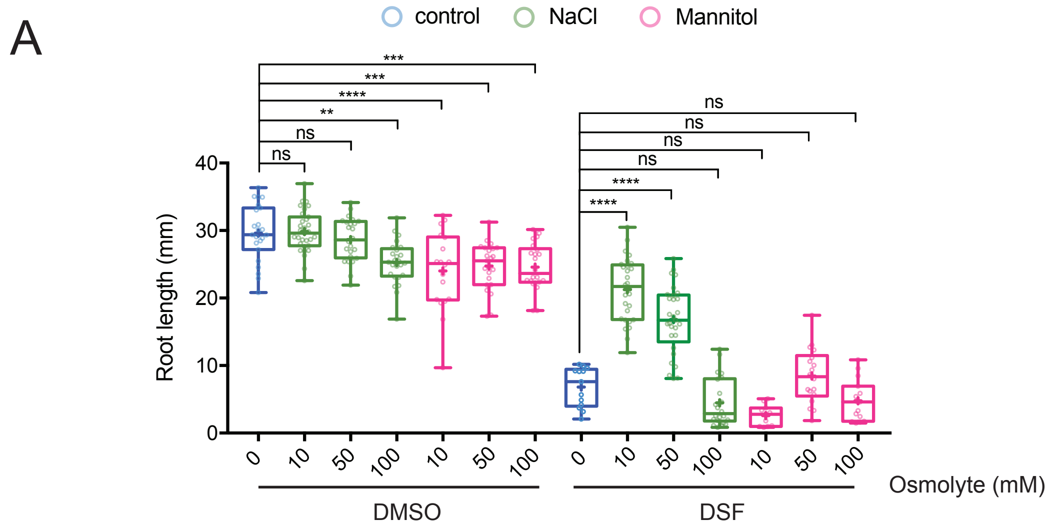
A
control
NaCl
Mannitol
***
***
****
**
ns
ns
ns
ns
ns
ns
****
****
Root length (mm)
40
30
20
10
0
0
10
50
100
10
50
100
0
10
50
100
10
50
100
Osmolyte (mM)
DMSO
DSF

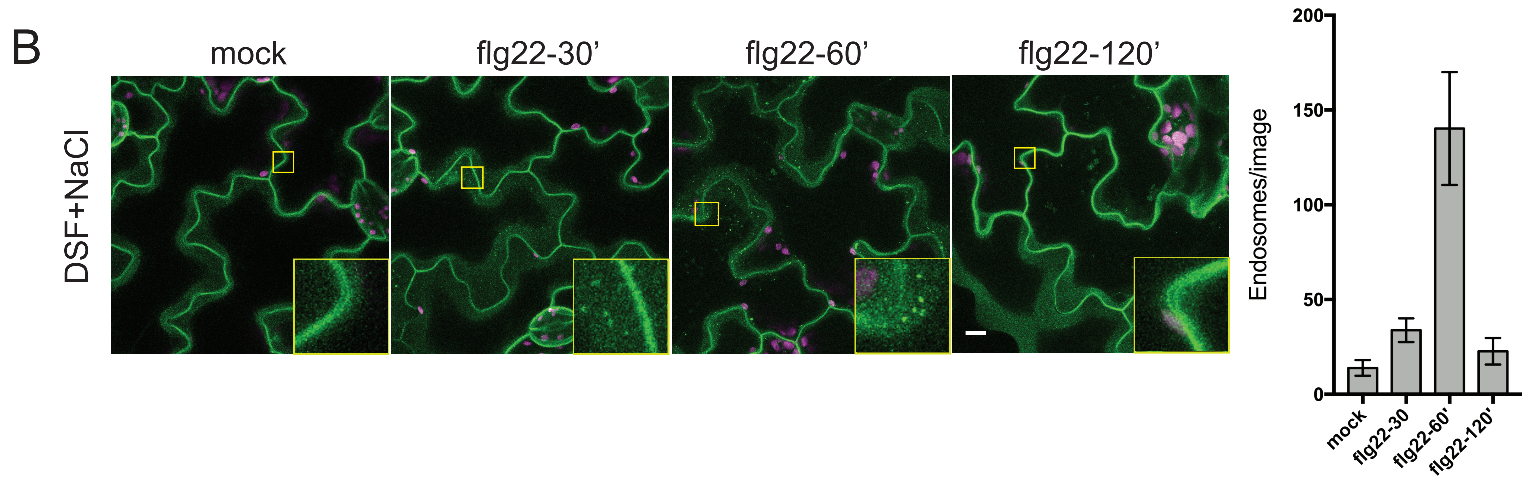
B
mock
flg22-30'
flg22-60'
flg22-120'
DSF+NaCl
Endosomes/image
200
150
100
50
0
mock
flg22-30
flg22-60'
flg22-120'

**Table S1. Key resources table**

| REAGENT or RESOURCE | SOURCE | IDENTIFIER |
|---|---|---|
| **Antibodies** | | |
| Rabbit Polyclonal anti-GFP | Torrey Pines BioLabs | Cat#TP401 |
| Rabbit Monoclonal anti-Phospho-p44/42-MAPK | Cell Signalling Technology | Cat#4370S |
| Mouse anti-FLAG M2 antibody | Sigma | Cat#A8592 |
| Rabbit anti-HA (3F10) antibody | Roche | Cat#12013819001 |
| Mouse anti-FLAG M2 affinity gel | Sigma | Cat#A2220 |
| **Bacterial and Virus Strains** | | |
| *Pseudomonas syringae* pv. *tomato* (*Pst*) DC3000 | 1 | N/A |
| **Chemicals, Peptides, and Recombinant Proteins** | | |
| *cis*-11-methyl-dodecenoic acid (DSF) | Sigma-Aldrich | Cat#42052 |
| ES9-17 | Chembridge | Cat#7577817 |
| Brefeldin A (BFA) | MPBio | Cat#159027 |
| Wortmannin | Sigma-Aldrich | Cat#W1628 |
| Lovastatin | Santa Cruz Biotech | Cat#SC-200850 |
| Methyl-$\beta$-cyclodextrin (M$\beta$CD) | MedChemExpress | Cat#HY-101461 |
| $\beta$–Sitosterol | Abcam | Cat#ab143122 |
| Stigmasterol | Santa Cruz Biotech | Cat#SC-281156 |
| Campesterol | Santa Cruz Biotech | Cat#SC-214658 |
| flg22 (QRLSTGSRINSAKDDAAGLQIA) | GL Biochem Ltd. | N/A |
| flgII-28 (ESTNILQRMRELAVQSRNDSNSATDREA) | GL Biochem Ltd. | N/A |
| **Critical Commercial Assays** | | |
| Rneasy Plant Minikit | Qiagen | Cat#74904 |
| RNAClean XP Kit | Agencourt | Cat#A63987 |
| SuperScript III First-strand synthesis system | Invitrogen | Cat#18080051 |
| Kapa SYBR FAST qPCR Master Mix (2X) Universal | Kapa Biosystems | Cat#KK4601 |
| Spurr Low-viscosity embedding kit | Sigma-Aldrich | Cat#EM0300 |
| **Experimental Models: Cell Lines** | | |
| Hamster: CHO cells: GFP-GPI | 2 | N/A |
| **Experimental Models: Organisms/Strains** | | |
| *Arabidopsis:* Col-0 | ABRC | CS1092 |
| *Arabidopsis:* RabF2b-YFP (Wave_2Y) | ABRC | CS781647 |
| *Arabidopsis:* Got1p homolog-YFP (Wave_18Y) | ABRC | CS781655 |
| *Arabidopsis*: FLS2-GFP | 3 | N/A |
| *Arabidopsis*: *pREM1.2::YFP:REM1.2* | 4 | N/A |
| *Arabidopsis*: Ler | 5 | N/A |
| *Arabidopsis*: *fkX-224* | 5 | N/A |
| *Arabidopsis*: *Ws* | 6 | N/A |
| *Arabidopsis*: *smt1-1* | 6 | N/A |
| *Arabidopsis*: BOR1-GFP | 7 | N/A |
| *Arabidopsis:* BRI1-GFP | 8 | N/A |
| *Arabidopsis:* CLC-GFP | 9 | N/A |
| **Oligonucleotides** | | |
| *EF1$\alpha$* F: TGAGCACGCTCTTCTTGCTTTCA | 10 | N/A |
| *EF1$\alpha$* R: GGTGGTGGCATCCATCTTGTTACA | 10 | N/A |
| *FRK1F:* ATCTTCGCTTGGAGCTTCTC | 11 | N/A |

| | | |
|---|---|---|
| *FRK1R:* TGCAGCGCAAGGACTAGAG | 11 | N/A |
| ***Plasmids*** | | |
| pHBT-FLS2-HA | 12 | N/A |
| pHBT-BAK1-FLAG | 12 | N/A |
| **Software and Algorithms** | | |
| Matlab | MathWorks | RRIS:SCR_001622 |
| Imaris | Bitplane | RRID:SCR_007370 |
| Fiji (ImageJ) | NIH | RRID:SCR_002285 |
| Graphpad PRISM | Graphpad | Ver. 7.0, RRID:SCR_002798 |
| Huygens | Scientific Volume Imaging | RRID:SCR_014237 |
| **Other** | | |
| Tecan plate reader | TECAN LifeSciences | Infinite M200Pro |
| AB StepOnePlus Real-Time PCR system | Applied Biosystem | Cat#4376600 |
